# Cortico-Subcortical Interactions in Overlapping Communities of Edge Functional Connectivity

**DOI:** 10.1101/2021.10.19.465016

**Authors:** Evgeny J Chumin, Joshua Faskowitz, Farnaz Zamani Esfahlani, Youngheun Jo, Haily Merritt, Jacob Tanner, Sarah A. Cutts, Maria Pope, Richard Betzel, Olaf Sporns

## Abstract

Both cortical and subcortical regions can be functionally organized into networks. Regions of the basal ganglia are extensively interconnected with the cortex via reciprocal connections that relay and modulate cortical function. Here we employ an edge-centric approach, which computes co-fluctuations among region pairs in a network to investigate the role and interaction of subcortical regions with cortical systems. By clustering edges into communities, we show that cortical systems and subcortical regions couple via multiple edge communities, with hippocampus and amygdala having a distinct pattern from striatum and thalamus. We show that the edge community structure of cortical networks is highly similar to one obtained from cortical nodes when the subcortex is present in the network. Additionally, we show that the edge community profile of both cortical and subcortical nodes can be estimates solely from cortico-subcortical interactions. Finally, we used a motif analysis focusing on edge community triads where a subcortical region coupled to two cortical regions and found that two community triads where one community couples the subcortex to the cortex were overrepresented. In summary, our results show organized coupling of the subcortex to the cortex that may play a role in cortical organization of primary sensorimotor/attention and heteromodal systems and puts forth the motif analysis of edge community triads as a promising method for investigation of communication patterns in networks.

## 1 Introduction

Characterizing the structural organization of the human brain has been a goal of network neuroscience since the inception of the connectome (Hagmann et al., 2008; Olaf Sporns, Tononi, & Kötter, 2005). The anatomical organization of connected neural elements is highly complex, which gives rise to a large repertoire of functional interactions. By aggregating blood oxygen level dependent (BOLD) signal from functional magnetic resonance imaging (fMRI) into regional measurements, functional connectivity (FC) among these regions (network nodes) can be estimated via measures of statistical dependence (network edges/weights; commonly Pearson correlation) between BOLD time courses. Resting state and task fMRI paradigms have revealed an intrinsic organization into functional systems (Fox, Corbetta, Snyder, Vincent, & Raichle, 2006; Yeo et al., 2011), which, as network science approaches have revealed, are arranged into a multi-scale modular architecture (Betzel & Bassett, 2017; Mucha, Richardson, Macon, Porter, & Onnela, 2010; O. Sporns & Betzel, 2016). Additionally, within clinical network neuroscience alterations in FC have been reported in various disease states such as schizophrenia (Fornito, Zalesky, Pantelis, & Bullmore, 2012), Alzheimer’s disease (Contreras et al., 2019), and other brain disorders (Fornito, Zalesky, & Breakspear, 2015).

The modular structure of fMRI-derived FC is a topic of great interest in network neuroscience. It refers to the decomposability of FC into clusters or communities of neural elements that possess greater connectivity within the same community compared to between different communities (Power et al., 2011). The brain’s community structure has been studied in relation to networks estimated from task fMRI reported activations (Stanley, Dagenbach, Lyday, Burdette, & Laurienti, 2014) and from intrinsic functional systems identified at rest (Betzel, Fukushima, He, Zuo, & Sporns, 2016; Di & Biswal, 2015). It is now hypothesized that modular structure is important for specialized brain function (Bertolero, Yeo, & D’Esposito, 2015). This hypothesis is supported by studies that have shown correspondence between intrinsic networks at rest and activations in task-based paradigms (Crossley et al., 2013; Di, Gohel, Kim, & Biswal, 2013), as well as reconfiguration of modular networks between rest and task (Cohen & Esposito, 2016; Hearne, Cocchi, Zalesky, & Mattingley, 2017; S. M. Smith et al., 2009). However, the focus has remained primarily on cortical community structure, with less work focused on contributions from major subcortical regions (Bell & Shine, 2016).

The basal ganglia and related structures within the subcortex are involved in a diverse set of functions though intra-subcortical communication as well as with cortical regions. Core regions of the subcortex, the striatum and globus pallidus are predominantly associated with motor functions and reward (Haber & Knutson, 2010; Lanciego, Luquin, & Obeso, 2012), while other subcortical structures (thalamus, amygdala, hippocampus) have roles in emotional, memory, and sensorimotor functions through various subcortical and cortico-subcortical circuits (Child & Benarroch, 2013; Choi, Ding, & Haber, 2017; Janak & Tye, 2015; Nakajima & Halassa, 2017; Sherman, 2017). Early understanding of subcortical function comes from studies that employed electrical stimulation and tract tracing in nonhuman models (Haber, Lynd, Klein, & Groenewegen, 1990; Olds & Milner, 1954), as well as lesion studies of human patients (Ward, Seri and, & Cavanna, 2013). As fMRI has become increasingly used to map subcortical organization and function noninvasively *in vivo,* studies that focused on subcortical FC have shown connectivity between the amygdala nuclei and hippocampus, caudate, and several cortical regions (prefrontal cortex, insula, and cingulate) (Janak & Tye, 2015; Tillman et al., 2018; Weis, Huggins, Bennett, Parisi, & Larson, 2019). Differential FC profiles of thalamic nuclei connectivity to cortical and subcortical regions have also been observed (Child & Benarroch, 2013; Nakajima & Halassa, 2017; Sherman, 2017). In recent years, methods have emerged that focus on network connections/edges more so than on network nodes (for review see (Faskowitz, Betzel, & Sporns, 2021)). Among those, Faskowitz, Esfahlani, Jo, Sporns, and Betzel (2020) developed a framework that represents the network as functional interactions of edges, which can be clustered to reveal an edge community structure, where each edge is assigned a community label (Faskowitz et al., 2020; Jo et al., 2020). This edge community structure approach offers a novel avenue for investigating subcortico-cortical interactions and communication.

The edge FC model uses nodal BOLD time series to estimate co-fluctuations among pairs of nodes, which can be interpreted as time-dependent pattens of communication. Therefore, community structure obtained from these co-fluctuation edge time series identifies groups of edges that may support similar communication strategies among connecting nodes and, when mapped back onto a node-by-node matrix, reveals an overlapping community structure. Prior work (Faskowitz et al., 2020; Jo et al., 2020; Zamani Esfahlani et al., 2020) has focused on mapping cortical communities, revealing a distinction between primary systems (visual, somatomotor, dorsal and ventral attention, and temporal parietal) and heteromodal systems (control, default mode, and limbic), without considering the roles and contributions of subcortical nodes. To address this, we investigated how subcortical regions contribute to, affect, or are affected by cortical modular organization. Our hypothesis is that the subcortex, via edge community structure, will differentially interact with different cortical systems. We also introduce a novel motif analysis based on edges’ community assignments, which we refer to as “edge community triads”, and we leverage this approach to further probe subcortico-cortical communication patterns. Edge community triads are three node subgraphs consisting of four types, based on the organization of edge communities. Given to the role of the subcortex in information integration, we hypothesized that triads which connect a subcortical node to two cortical nodes via the same edge community will be over-represented relative to other triad types. This would demonstrate that subcortico-cortical communication patterns as estimated by edge community structure from resting state fMRI (rs-fMRI) capture biologically meaningful information about subcortico-cortical organization.

## 2 Materials and Methods

### 2.1 Dataset

In this study we analyzed data from the Human Connectome Project (HCP) ICA-FIX (Griffanti et al., 2014; Van Essen et al., 2013) preprocessed dataset. Informed consent was obtained from all participants and all study protocols and procedures were approved by the Washington University Institutional Review Board. Data were collected on a Siemens 3T Connectom Skyra with a 32-channel head coil. A detailed description of acquisition protocols can be found elsewhere (Glasser et al., 2013; Van Essen et al., 2013). Briefly, rs-fMRI data were acquired in 4 sessions over 2 days, (scan duration 14:33 min) with a gradientecho echo-planar imaging sequence, with TR=720ms, TE=33.1ms, flip angle of 52 degrees, 2mm isotropic voxel resolution and a multiband factor of 8. Participants were instructed to keep eyes open and fixated on a cross. A subset of 92 unrelated participants, part of the HCP 100 unrelated subjects release, for which complete processed data was available for all four scans, were utilized in primary analyses. Scans were excluded from analysis based on a set of summary motion measurements (Parkes, Fulcher, Yücel, & Fornito, 2018) derived from motion correction preprocessing and provided in the HCP database. Motion spikes were defined as a relative root-mean-square (RMS) movement of 0.25mm or above. Scans were excluded from analysis if at least one of the following conditions were met: greater than 15% of time points were marked as a motion spikes; the average relative RMS motion greater than 0.2mm; a spike larger than 5mm was present. These criteria resulted in eight subjects with an incomplete set of fMRI scans, and therefore excluded from the present study.

### 2.2 Image pre-processing

#### Functional preprocessing

HCP rs-fMRI data were minimally preprocessed as described in Glasser et al. (2013) including distortion, susceptibility, and motion correction, registration to subjects’ respective T1-weighted data, bias and intensity normalized (mean 10,000), projected onto the 32k_fs_LR mesh, and aligned to common space with a multi-modal surface registration (Robinson et al., 2014). In addition to the ICA-FIX artifact removal process, global signal, its derivative, and their squared terms were regressed out, and data were detrended and bandpass filtered (0.008 – 0.08 Hz) (Parkes et al., 2018) with Nilearn signal.clean, which removes confounds orthogonal to the temporal filters (Lindquist, Geuter, Wager, & Caffo, 2019).

#### Parcellation pre-processing

A functional parcellation of the cortex, designed to optimize both local gradient and global similarity measures of fMRI signal was used to define nodes at 4 scales (Schaefer100-400 nodes in steps of 100) (Schaefer et al., 2018). This parcellation is mapped to and grouped by canonical 17 resting state networks from Yeo et al. (2011). Parcellations were downloaded as *cifti* files in the *fsLR_32k* space; the same space as the preprocessed rs-fMRI data. Additionally, a novel gradient-based subcortical parcellation was used to delineate nodes within the amygdala, hippocampus, thalamus, and striatum consistent of 16 regions per hemisphere (Scale II parcellation from Tian, Margulies, Breakspear, and Zalesky (2020)).

### 2.3 Edge graph construction

Mean nodal time courses were extracted from preprocessed data for each scan and each cortical and subcortical parcellation. For each subject, time courses were cropped to exclude the first and last 50 timepoints (~36 seconds) to account for the edge effects of the bandpass filter and concatenated into day 1 and day 2 (2 scans each). Subsequently, sample time courses were constructed for each day by concatenating all subject time courses from that day.

Edge time series (ETS) were computed as described previously (Faskowitz et al., 2020; Jo et al., 2020), in the following steps: (1) for all nodes (*N*), time courses were *z*-scored, (2) for all possible node pairs *i* and *j* (where *i* ≠ *j),* the element-wise product (over time) was computed. The resultant 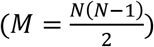 ETS are vectors of same length as the nodal time courses that encode moment-by-moment co-fluctuations of the edge between nodes *i* and *j.* ETS can be interpreted as the decomposition of FC (Pearson correlation) into its time-varying contributions.

### 2.4 Edge community detection

Computing a matrix from ETS would result in large (*M x M)* matrices that would require a great amount of memory and computation time to cluster. To circumvent this issue and reduce computational burden, we clustered the *M* × *T* ETS matrix directly using the k-means algorithm, implemented in MATLAB version 2021A, with normalized Euclidean distance (Jo et al., 2020). We varied the number of clusters, *k,* from 2 to 20 clusters in increments of 1, repeating the algorithm 250 times with random initial conditions. Due to the large number of time points, for sample representative communities each run was initiated with 10% of the concatenated time series randomly sampled and clustered to produce an initial estimate of cluster centroids. These centroids were then used as initial estimates to cluster the full sample time series. At each k value a single consensus partition was obtained from the 250 runs. For subject-specific partitions, the clustering algorithm was run (1 run) with the sample consensus partition provided as the initial seed partition. Note that no additional runs were necessary, as the k-means algorithm is deterministic; from a fixed initial assignment of nodes to clusters it always converges to the same solution.

### 2.5 Community overlap metrics

#### Normalized Entropy

Interpreted as a continuous measure of edge community overlap at any node *i*, normalized entropy was calculated by first computing node *i’s* participation in cluster *c*:

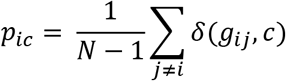

where *g_ij_* ∈ (1, …, *k*) was the cluster assignment of the edge between nodes *i* and *j* and *δ*(*g_ij_, c*) is the Kronecker delta, whose value is 1 if *x* == *y* and zero otherwise. The entropy of the probability distribution *p_i_*, = [*p*_*i*1_,…, *p_ik_*] of node *i’s* edge community assignment was then computed as:

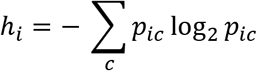

This value was then normalized by dividing by log_2_ *k,* which bounded the range to [0,1]. For sample time series derived clusters, entropy is reported as the mean across the 250 repetitions of the clustering algorithms.

#### Edge community similarity

The resultant partition from the clustered ETS is a vector where each edge is assigned a community label. This vector can be rearranged into the upper triangular of a *N * N* matrix *X*, where row/column for any node *i* encodes the community affiliations for edges connecting that node. Similarity of edge communities can then be computed from this matrix for nodes *i* and *j* as the fraction of edges that have the same community labels for both nodes (Jo et al., 2020):

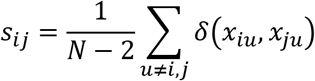

where *δ*(*x_iu_, x_ju_*) is the Kronecker delta that is 1 if x and y have the same value and 0 otherwise. Repeating the process over all node pairs generates the similarity matrix S.

#### Edge community triads

A triad of nodes can be analyzed as a three-node motif, a mesoscale building block of the wider brain network (Milo et al., 2002). Motifs allow for a complete decomposition of a larger network into subgraphs, which can reveal statistical features in the local organization of structural and functional brain networks (Battiston, Nicosia, Chavez, & Latora, 2017; Olaf Sporns & Kötter, 2004). Here, we adapt graph-based motif analysis to include edge labels based on their edge community assignment, which allows for investigation of relationships among nodes using triad motifs in a fully connected network. Focusing on a reference node (indexed *l* in Fig. 1) and examining all or subset of triads it takes part in allows us to ask questions about communication/coupling properties of a that node to other nodes of the network, based on edge communities that make up the triads. Here the focus is on cortico-subcortical connectivity, thus we focus our analysis on triads with a single subcortical node in the reference position and examine the coupling of these reference nodes to all possible pairs of cortical nodes.

**Figure 1.**
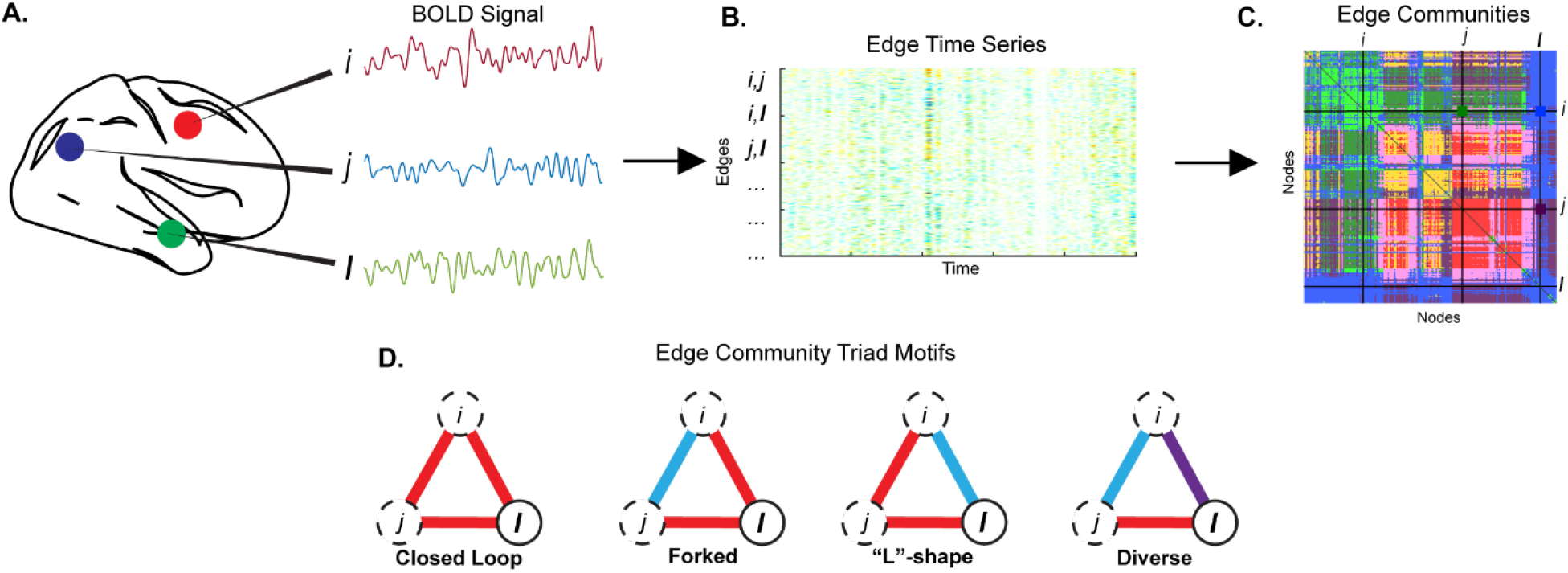
Edge time series clustering overview. Nodal time series **(A)** for all regions pairs are used to compute the edge time series **(B)**, which are subsequently clustered to assign each edge in the network to an edge community, as visualized in the node-by-node matrix **(C)**. From this matrix motif triads are estimated, with four possible triad types **(D)**: a closed loop comprised of a single community, a forked triad (relative to a reference node *l*) comprised of two communities with the same community for both edges connected to *l,* L-shape triad also comprised of two communities, however, edges connecting *l* are in different communities, and a diverse triad comprised of three different communities.

Figure 1 shows a diagrammatic workflow from nodal time series to edge community triads. There are four possible edge community triads, in any network where the number of communities is greater than two: (1) closed loop – the three edges among the three nodes are all in the same community, (2) forked – the two edges connecting to/from the reference (subcortical) node have the same community label, while the third edge (between the two cortical nodes) is in a different community, (3) L-triad – the two edges connecting to/from the reference node have different community labels, while the third edge (that does not connect the reference node) shares a label with one of the two reference connecting edges, and (4) diverse – all three edges in the triad have different community labels. A diagram of the possible triads is shown in Fig. 1D. To assess whether the distributions of triad types in a network are different from those expected by chance, a set of 1,000 null networks was generated from the node-by-node edge community assignment matrix, permuting the node labels in such a way that the indices of the subcortex were permuted into the cortex portion of the matrix. This null was chosen over a blind label permutation to ensure that anything that could make the subcortex distinct from the cortex in terms of its triad pattern was restructured under the null. This null distribution was used to compute a permutation *p*-value.

## 3 Results

### 3.1 Consensus edge community structure

Here we investigated the community structure of sample concatenated time series, using a parcellation of 200 cortical and 32 subcortical nodes, by performing edge-centric clustering of concatenated time series of 92 participants from the HCP cohort. Clustering of the edge time series leads to community partitions where each edge is assigned a label and these partitions can be projected onto a node-by-node matrix to visualize overlapping community structure (Fig.2A). This approach has been previously applied to cortical brain networks (Faskowitz et al., 2020; Jo et al., 2020; Zamani Esfahlani et al., 2020) and we extend this work by assessing the role/influence of subcortical regions in the brain.

**Figure 2.**
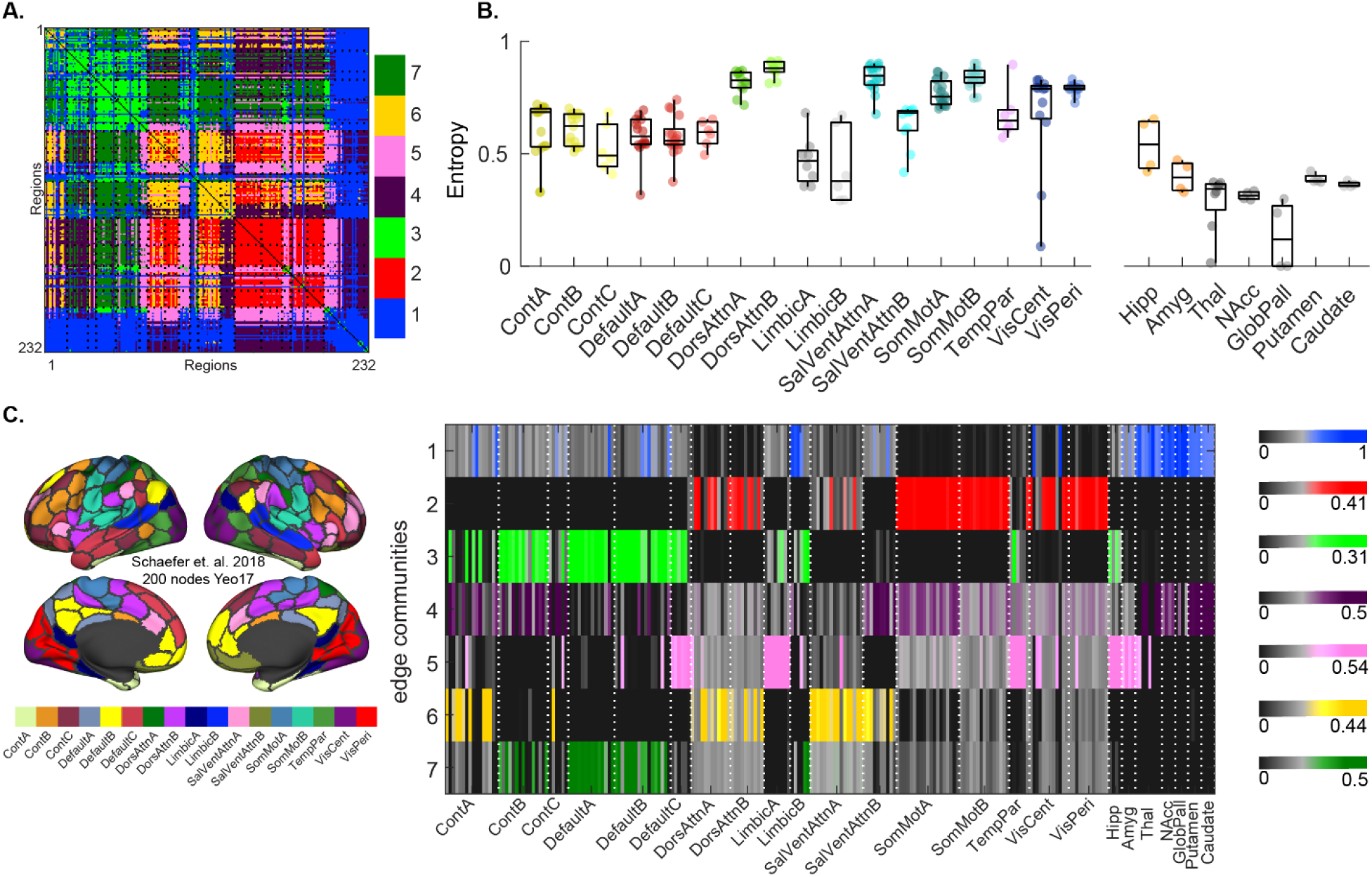
**(A)** Edge community affiliations for the 7-cluster solution represented in a node-by-node matrix for the 232 node parcellation (200 cortical Schaefer et al. (2018) nodes and 32 subcortical Tian et al. (2020) nodes). Black dotted lines denote boundaries of 17 resting state networks from Yeo et al. (2011) plus subcortical. Networks are ordered top-bottom and left-right as labeled on x-axis in B and C. **(B)** Nodal entropies of edge communities grouped into 17 networks plus subcortical nodes, which are grouped by anatomy. Individual data points denote node groupings into the 17 canonical RSNs in Yeo et al. (2011), colored by 7 network labels, with added subcortical regions. **(C)** (Left) A surface template of the Schaefer et. al., 2018 node assignment to the Yeo et. al., 2011 17-networks. (Right) Edge communities (y-axis) across resting state networks and subcortex (x-axis). Color saturation corresponds to proportion of total edges in an edge community that connect to a node of a particular network. Variants of these plots for different cluster solutions, cortical parcellation scales, and for the Day 2 dataset are in Supplementary Figs. 1–3. Cont – Control, DorsAttn – Dorsal Attention, SalVentAttn – Salience/Ventral Attention, SomMot – Somatomotor, TempPar – Temporal Parietal, VisCent – Visual Central, VisPeri – Visual Peripheral, Hipp – Hippocampus, Amyg – Amygdala, Thal – Thalamus, NAcc – Nucleus Accumbens, GlobPall – Globus Pallidus.

When ordered by canonical RSNs, estimated partitions showed overlapping organization that qualitatively resembled canonical RSN grouping, with a distinct subcortical component (Fig. 2A). The distribution of edge communities for all nodes was computed as normalized entropy and visualized by system for cortical RSNs and subcortical nodes, which were grouped by anatomical label (Fig. 2B), showing a potential dichotomy of primary sensory and attentional systems (visual, somatomotor, dorsal and ventral attention, and temporal parietal) in one group and higher order systems (control, default mode, and limbic) in another, as previously reported (Jo et al., 2020). A comparison of mean network entropies among the two system types showed significantly higher entropy values were observed in primary sensorimotor and attentional systems (mean ± standard deviation: 0.7673 ± 0.0877), indicating that edges incident upon nodes of those networks are distributed over a greater number of communities, compared to higher order systems (0.5490 ± 0.0630) (permutation *t*-test, *p* = 0.00011, 100,000 label permutations). Similar outcomes were observed at other cluster solutions (number of clusters = 4, 10, and 17; Supplementary Fig.1), with a range of cortical parcellation scales (Schaefer 100, 300, and 400 nodes; Supplementary Fig. 2), and in a second dataset consisting of Day 2 HCP scans from the same participants (Supplementary Fig. 3).

Contributions of edge communities to RSN systems as well as the contribution of subcortical nodes is shown in Figure 2C. For the 7-cluster solution, three communities were predominantly cortical (communities 2,6,7) and coupled visual-somatomotor-attention, attention-control, and control-default mode systems, respectively. Four edge communities coupled subcortical nodes to cortical systems, which qualitatively showed two subcortical groupings (Fig. 2C, communities 1 and 4: striatum, pallidum, and thalamus; communities 3 and 5: predominantly hippocampus and amygdala) coupled to either the primary sensorimotor/attention or heteromodal systems. Anatomical visualizations of edge distributions for each of the seven communities at each node are shown in Figure 3 as well as in Supplementary Fig.4 for the Day 2 HCP dataset, where spatially similar communities were observed.

**Figure 3.**
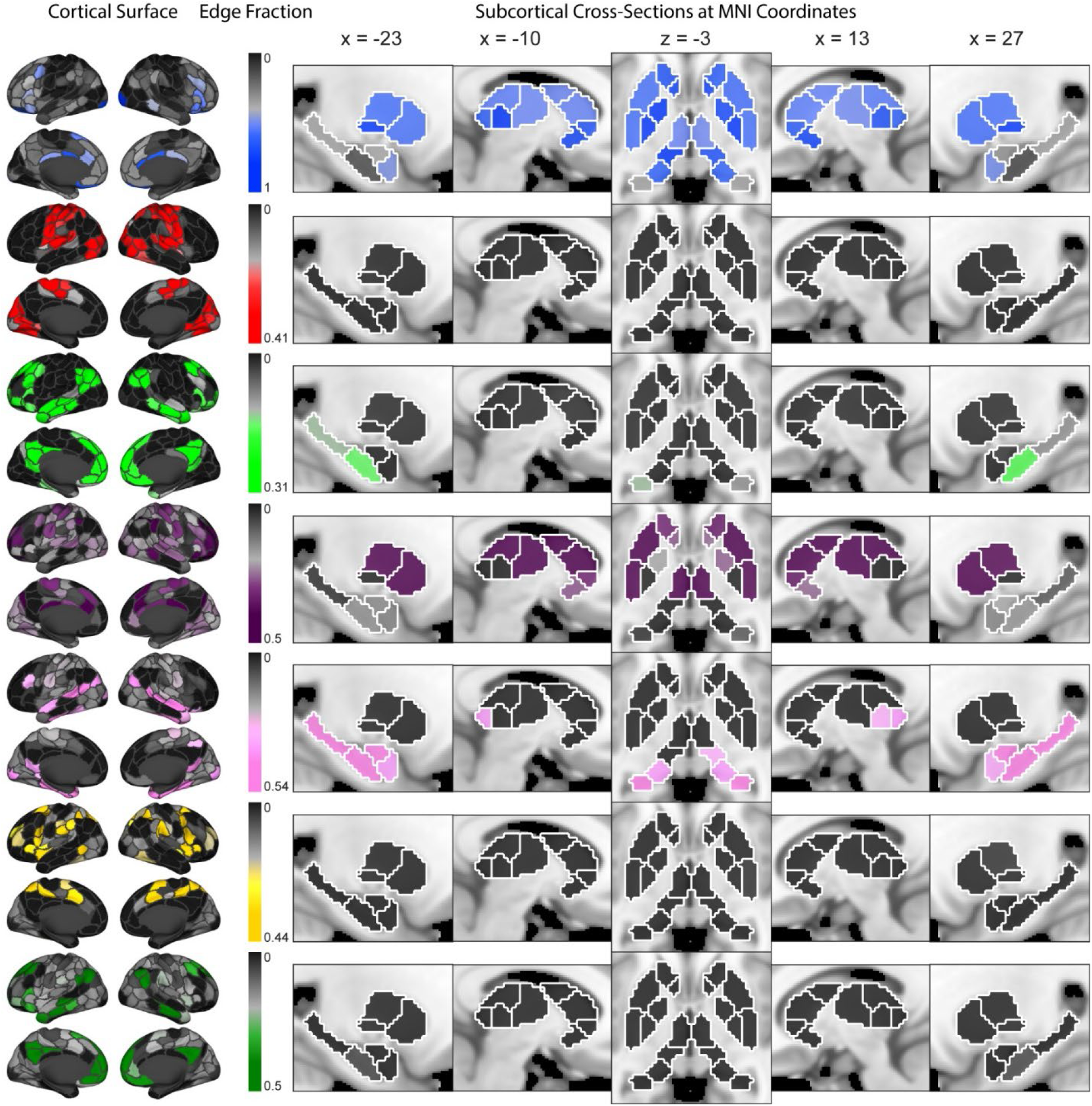
Spatial representation of edge communities for cortical (Left; 200 nodes from Schaefer et al. (2018)) and subcortical (Right; 32 nodes from Tian et al. (2020)) nodes. Color saturation corresponds to proportion of total edges in an edge community that connect to a particular node. Cortical nodes are visualized on a fs_LR_32k surface, while subcortical nodes are sagittal slices and an axial slice in Montreal Neurological Institute (MNI) standard space at the indicated coordinates. Anatomical underlay is the MNI152_1mm brain template. Slices left of the axial slice are in the left hemisphere while slices to the right are in the right hemisphere.

### 3.2 Role/Influence of subcortical nodes on cortical edge community structure

Given previous reports of edge community organization within cortical systems (Jo et al., 2020; Zamani Esfahlani et al., 2020), how does the presence of subcortical nodes impact edge community structure? We assessed this by comparing cortical edge communities for the 7-cluster solutions for the full network (cortical + subcortical) vs. cortex only (Fig. 4). Edge communities for cortical nodes were highly similar between the two networks (normalized mutual information 0.7153; Fig. 4A) and comparison across a range of community solutions showed high similarity for communities with similar number of clusters (Fig. 4B). Similar separation of primary and heteromodal systems via edge communities (Fig. 4C) and similar system entropies (Fig. 4D; primary 0.7293 ± 0.1009 and heteromodal 0.5175 ± 0.0784 systems, permutation *t*-test, *p* = 0.0004, 100,000 label permutations) were also observed. To further examine how edge communities are related to nodal RSN groupings, the influence of subcortical nodes, and the distinction between primary and heteromodal RSNs, edge community profile similarity matrices, which quantify the degree to which two nodes couple to other nodes in the network via the same edge communities, were generated. Comparing the edge community similarity of cortical nodes showed that addition of subcortical nodes to the network had minimal influence on similarity (Fig. 4E-F, Pearson correlation r = 0.94 across cortical nodes between two network types) This relationship was also observed at varying numbers of clusters and at other cortical parcellation scales (Supplementary Fig. 5) as well as in the day 2 dataset (Supplementary Fig. 6).

**Figure 4.**
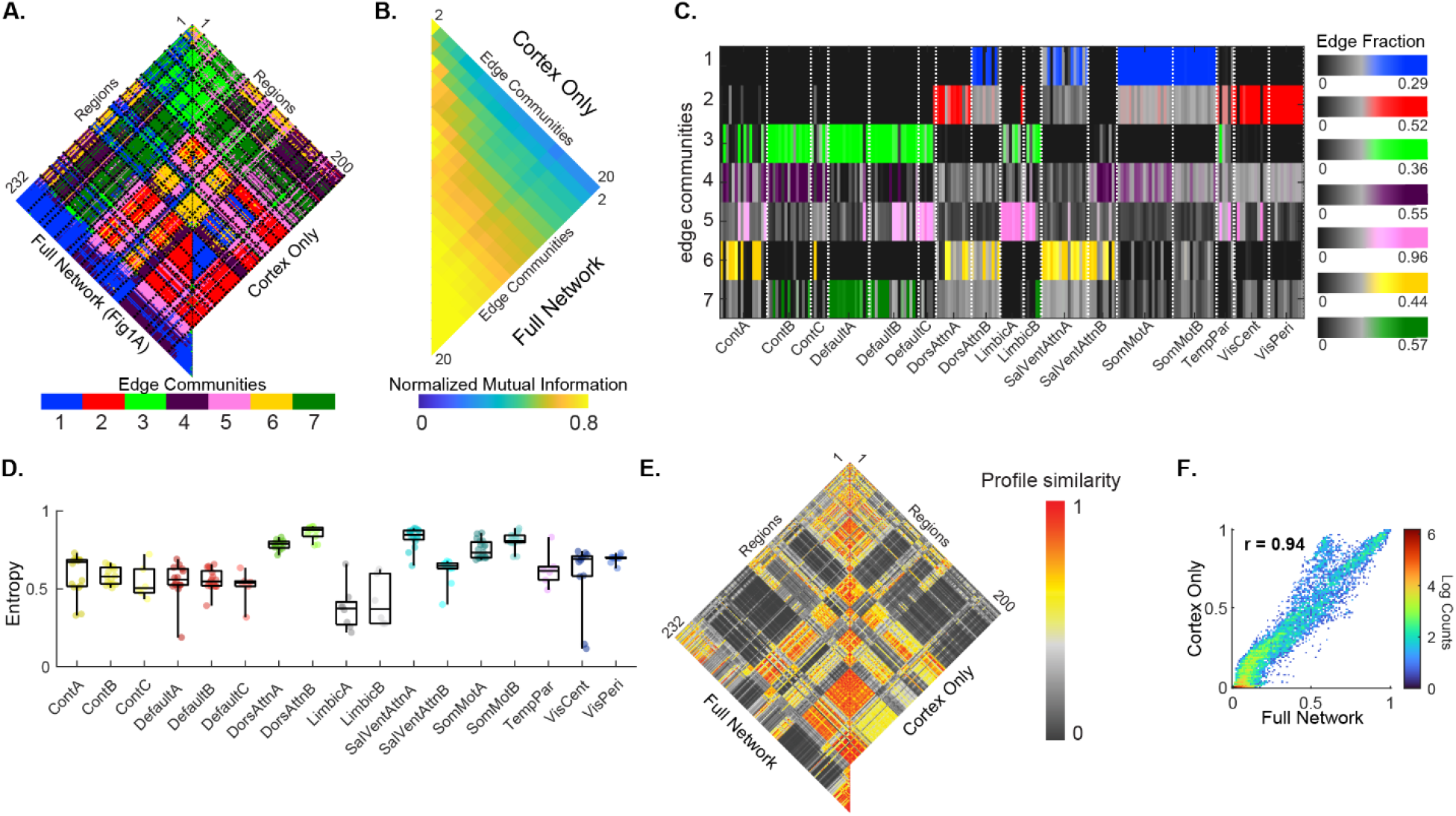
Cortico-cortical edge community structure is preserved with addition of subcortex. **(A)** Edge community clusters of the full (cortex + subcortex, as seen in Figure 2) network (Left) compared to when only cortical edges were clustered (Right). Black dotted lines denote boundaries of the Yeo et al. (2011) 17 resting state networks. **(B)** Similarity of edge community partitions of varying sized (from 2 through 20) for cortical nodes from the cortex only network compared to cortical nodes from the cortex + subcortex network. **(C)** Overlapping edge communities (y-axis) across resting state networks (x-axis) of the cortical nodes only network. Color saturation corresponds to proportion of total edges in an edge community that connect to a node of a particular network. **(D)** Nodal entropies of edge communities grouped by 17 networks in the cortex only matrix, with individual data point colors denoting groupings into the 7 canonical RSNs in Yeo et al. (2011). **(E)** Comparison of edge community profiles (similarity) of cortical nodes from the full network (left) versus the cortex only network (right). **(F)** Density plot of the edges from the two network types in D with Pearson correlation reported. Cont – Control, DorsAttn – Dorsal Attention, SalVentAttn – Salience/Ventral Attention, SomMot – Somatomotor, TempPar – Temporal Patietal, VisCent – Visual Central, VisPeri – Visual Peripheral, r – Pearson Correlation.

Focusing on the edge communities within the cortico-subcortical interaction block (Fig. 5A), the distribution of the interaction edges of communities with subcortical components among the RSNs shows a distinction between hippocampus/amygdala and striatum/thalamus edge community profiles. Nodal entropies computed only from the interactions between the cortex and subcortex (i.e., row entropy for subcortical nodes and column entropy for cortical nodes) recapitulate the pattern of segregation of primary (0.4557 ± 0.0729) and heteromodal (0.2105 ± 0.0956) systems (permutation *t*-test, *p* = 0.0002, 100,000 label permutations). Just as with entropy, edge community profile similarity can be computed only from the cortico-subcortical interaction elements. For subcortical nodes, profiles were nearly identical for the full network versus the interaction block (Fig. 5C). Cortical edge community profiles from the interaction block were also highly similar to profiles from the full network albeit more variable (Fig. 5D). The finding that edge community profiles from cortico-subcortical interactions reflect cortical and subcortical profiles obtained from the full network are consistent at various cluster solutions (Supplementary Fig. 7), a range of cortical parcellation scales (Supplementary Fig. 8), and in the Day 2 data from these participants (Supplementary Fig. 9).

**Figure 5.**
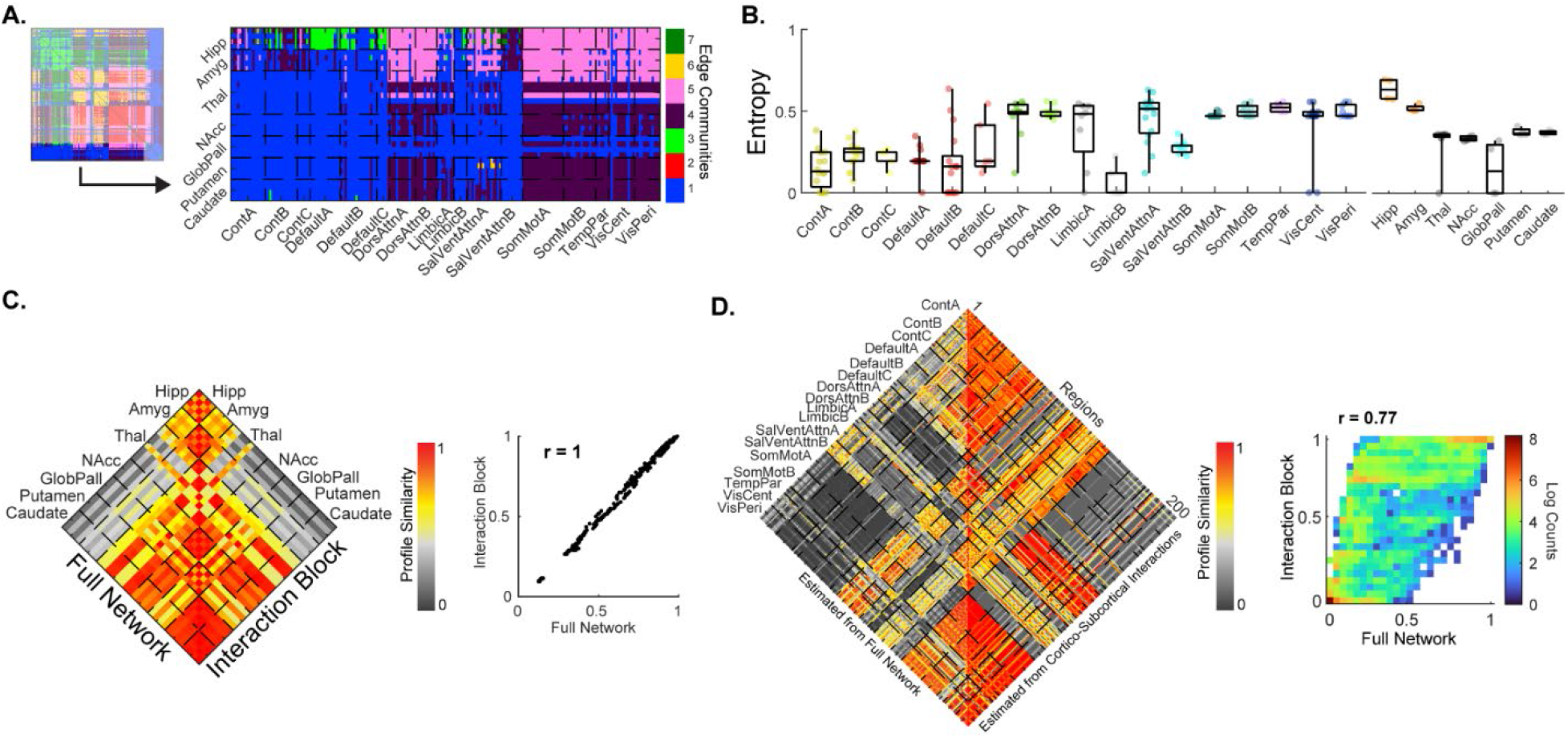
**(A)** Zoomed in view of edge community structure of cortico-subcortical interactions. Black dashed lines denote boundaries for either the subcortex, grouped by anatomy (y-axis), or the Yeo et al. (2011) 17 resting state networks (x-axis). **(B)** Cortical and subcortical node entropies computed only from interaction edges connecting to/from the subcortex and cortex, respectively. **(C)** Similarities computed from all edges (left; full network) and from edges within subcortex only (right; interaction block). Scatterplot shows edge to edge correlation between the similarities from the two network types. **(D)** Similarity among cortical nodes estimated from the full network (cortex + subcortex; left) and from only the subcortex interaction edges (right), with a 2D heatmap showing edge-to-edge correspondence of similarities from the two network types. Cont – Control, DorsAttn – Dorsal Attention, SalVentAttn – Salience/Ventral Attention, SomMot – Somatomotor, TempPar – Temporal Patietal, VisCent – Visual Central, VisPeri – Visual Peripheral, Hipp – Hippocampus, Amyg – Amygdala, Thal – Thalamus, NAcc – Nucleus Accumbens, GlobPall – Globus Pallidus, r – Pearson correlation.

### 3.3 Individual variability in edge community structure

While it is computationally infeasible to perform iterative clustering on edge time series from the 92 HCP participants individually, the consensus partition described in previous sections can be used as a seed/initial partition and clustering is then run once per participant to obtain an individual partition. Using this method, do the individual partitions recapitulate what was observed in consensus and what insight do they provide into cortico-subcortical coupling? Individual edge community partitions showed greater similarity to the consensus partition (median mutual information = 0.2984) than amongst each other (0.1765) (Fig. 6A), as expected. Entropies per RSN, computed per participant as the mean of the nodes in that system are shown in Fig. 6B. The median entropy for each participant across either primary, heteromodal, or subcortical systems is shown in Fig. 6C, where entropies were significantly different among the three system types (means ± standard deviations: primary 0.7894 ± 0.0477, heteromodal 0.6605 ± 0.0553, and subcortical 0.4839 ± 0.1069, permutation ANOVA, *p* < 0.00001, 100,000 label permutations; all post-hoc *t*-tests comparisons *p* < 0.001, Bonferroni adjusted), consistent with results obtained from consensus data. For edge community profile similarity, median values were extracted per participant for edges within primary, heteromodal, and subcortical nodes (Fig. 6D), as well as for edges between them (Fig. 6E). Profile similarities were significantly lower in heteromodal (0.3391 ± 0.0548) compared to primary (0.4444 ± 0.0738) or subcortical (0.4458 ± 0.1067) system types (Fig. 6D, permutation ANOVA *p* < 0.00001, 100,000 label permutations; post-hoc *t*-tests *p* < 0.001 Bonferroni adjusted, for heteromodal versus primary and subcortical system types). Finally, to probe whether the subcortical regions differed in their coupling to cortical systems, similarity values from edges coupling primary-heteromodal (0.1494 ± 0.0549), primary-subcortical (0.2006 ± 0.0549), and heteromodal-subcortical (0.2781 ± 0.0615) were compared. A permutation ANOVA with 100,000 label permutations showed a significant main effect of group (*p* < 0.00001), with the three groups significantly different from each other (Fig. 6E, all post-hoc *t*-tests *p*<0.001, Bonferroni adjusted). This pattern was consistent using different numbers of edge communities (Supplementary Fig. 10), at varying cortical parcellation scale (Supplementary Fig. 11), and in the Day 2 data (Supplementary Fig. 12).

**Figure 6.**
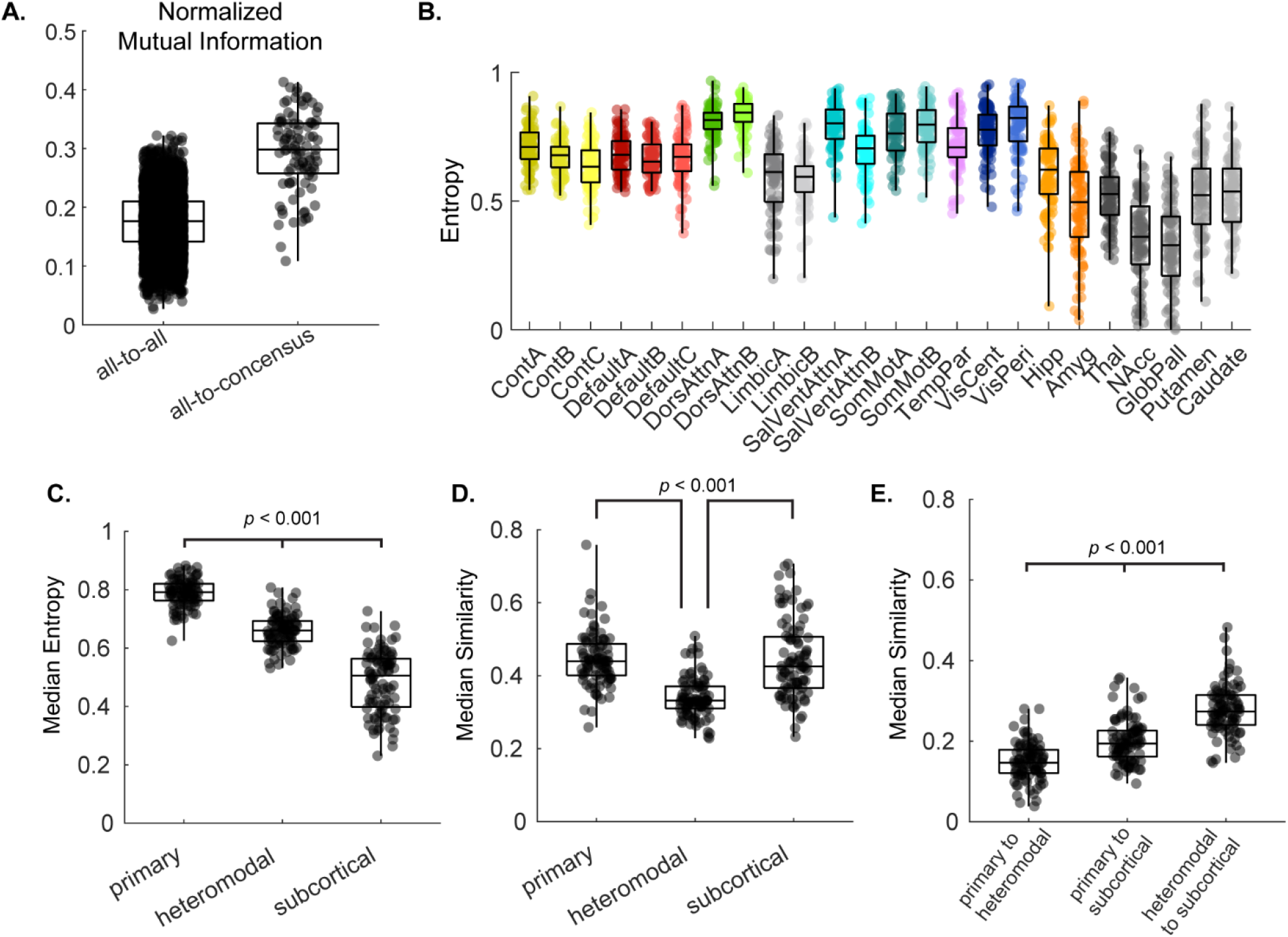
Assessment of variability and comparison to sample-wide data of individual subject-derived edge communities, with group consensus solution as initial condition. **(A)** Individual partition comparisons among all subject pairs (left) and of each subjects’ partition to the consensus partition from the sample concatenated time courses. **(B)** Nodal entropies of edge communities for individual subjects (each dot is a value for a node of a subject for a particular network) grouped by the 17 resting state systems and subcortical anatomical regions. Individual data point colors of cortical networks denote groupings into the 7 canonical RSNs in Yeo et al. (2011). **(C)** Median entropies computed across nodes in primary, heteromodal, and subcortical systems for each subject. **(D-E)** Median edge community profile similarities for subjects grouped as either **(D)** within system type or **(E)** between system types. Statistical significance denotes post-hoc Bonferroni adjusted *p*-value from an ANOVA. Cont – Control, DorsAttn – Dorsal Attention, SalVentAttn – Salience/Ventral Attention, SomMot – Somatomotor, TempPar – Temporal Patietal, VisCent – Visual Central, VisPeri – Visual Peripheral, Hipp – Hippocampus, Amyg – Amygdala, Thal – Thalamus, NAcc – Nucleus Accumbens, GlobPall – Globus Pallidus.

### 3.4 Subcortico-cortical communication via edge community triads

The above results show a dichotomy of canonical RSN systems into a primary sensorimotor and attention group and a higher order heteromodal group, which may be shaped by or influencing subcortical edge community organization. To further investigate subcortico-cortical coupling, we adapted the concept of motifs, which has been primarily used to quantify subgraphs in structural/sparse networks (Olaf Sporns & Kötter, 2004), focusing on triads (triangles of 3 nodes) that are comprised of one subcortical (reference node; Fig. 1D) and two cortical nodes. With each edge assigned a community label, four possible classes of triads can be identified: a closed loop, a forked triad, L-shape triad, or a diverse triad (Figs. 1D and 7A show a diagram of triad types). In this framework, how are the triad types distributed in instances of subcortico-cortical communication and what can this tell us about potential communication strategies between the cortex and subcortex?

**Figure 7.**
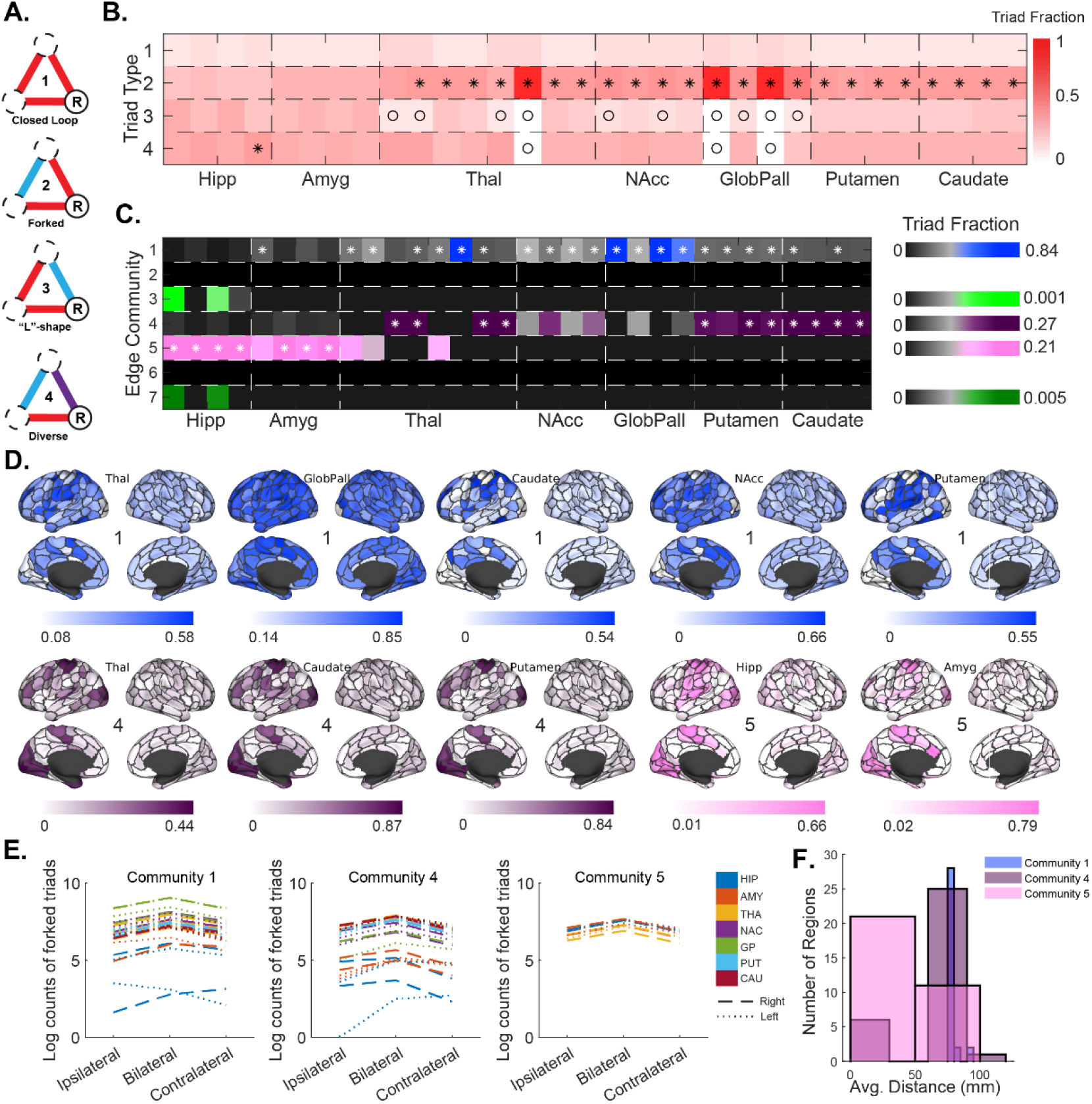
Edge community triads and their distribution around subcortical nodes. **(A)** Distribution of triads around subcortical nodes (x-axis) that connect it to any two cortical nodes. Rows correspond to triad types, from top to bottom the triads are closed-loop, forked, L-shape, and diverse. Columns sum to 1 (i.e., all triads that connect that subcortical node to cortical nodes). Compared to a null distribution of 1,000 networks, asterisks denote proportions that significantly exceed the null, while open circles indicate lower than null (permutation *p* < 0.05 two-tailed). **(B)** A breakdown of forked triads by edge community. Edge community assignments correspond to the community that connected to/from the subcortical reference node. Color saturation indicates proportions of all triads around the node. Asterisks denote a significantly greater fraction compared to 1,000 null networks (permutation *p* < 0.05, two-tailed). **(C)** For subcortical anatomical regions where more than half its nodes had significance in A, the cortical surface overlays show the average (across nodes within a subcortical region) fraction that a cortical node was part of the triad to that subcortical node. **(E)** Locations of cortical nodes relative to subcortical reference regions across triads. Colors denote subcortical anatomical labels, while dashed and dotted line denote right and left hemispheres, respectively. **(F)** Average distances (in mm) between cortical nodes for triads in each community, for all subcortical reference regions.

To answer this question, triad type distributions were generated for each subcortical node, connecting to two cortical nodes, from the original node-by-node edge community matrix as well as for 1,000 null matrices where node labels were permuted, with the condition that subcortical nodes did not remain in subcortex assigned indices after permutation. When compared to the null distribution, forked triads were present in a significantly higher fraction for the striatum, pallidum, and five of the six nodes of the thalamus (Fig. 7B; permutation *p* < 0.05, two-sided). Additionally, L-shape triads were present in lower frequency than expected for nodes of the thalamus, nucleus accumbens, and pallidum. In terms of communication strategies, forked triads imply that the reference (subcortical) node employs similar patterns of communication with each of the two cortical nodes, while these cortical nodes maintain a different pattern. This may point to a role of the subcortex in information integration and cortical modulation. L-triads, in turn, could be representative of a different (and statistically underrepresented) strategy of relay communication from subcortex to cortex or vice versa. Breaking down the forked triads by their edge community affiliation (the community that has two edges in the triad between subcortex and cortex), three communities (1, 4, and 5 in Fig. 7C) showed significantly higher fractions that expected by the null distribution (permutation *p* < 0.05). This finding is in line with the observation that large portions of these edge communities are in the subcortico-cortical interaction block (Fig. 5A). Visualization of the cortical nodes for these subcortico-cortical forked triads (Fig. 7D), qualitatively shows a distinction between edge community 1, which has distributed edges from the striatal/thalamic nodes to heteromodal systems and Salience/Ventral attention, and edge communities 4 and 5, which connect primary/attentional systems to striatum/thalamus and hippocampus/amygdala, respectively. Detailed matrices of cortical forked triad endpoints for each subcortical node are shown in Supplementary Figure 13. At other investigated number of communities (Supplementary Fig. 14) as well as varying cortical parcellations (Supplementary Fig. 15) similar outcomes were observed.

To better understand the cortical endpoints of forked triads, each triad was labeled as ipsilateral (same hemisphere relative to the subcortical reference node), bilateral (cortical endpoints in different hemispheres), or contralateral, with the counts for each subcortical node plotted in Fig. 7E. Overall the distributions were highly similar, with the anterior hippocampal nodes being the only notable standout (Fig. 7E blue lines), however, the triad counts for those nodes were orders of magnitude lower (tens vs hundreds/thousands). Therefore, the asymmetry observed in the anterior hippocampus is likely due to low counts. Next, to further understand the spatial distributions of forked triads over the cortex, the Euclidian distance between cortical nodes (in standard space) was computed and summarized as a distribution (for each edge community) of average distance between cortical nodes (for forked triads at each subcortical reference node). For the first edge community, where forked triads were overrepresented for thalamus and striatum, the distance between cortical nodes was ~100mm, which given the mostly equal distribution of ipsi-, bi-, and contralateral localization suggests they are coupling some combination of frontal and parietal nodes, either within or between hemispheres. This is consistent with Supplementary Fig. 13, which shows that the cortical endpoints fall in control, default, and ventral attention systems as they are primarily localized in frontal and parietal lobes. Edge community 4, which also involves the stratum and thalamus, showed similar length distributions, however, cortical endpoints of forked triads within this community fell in visual, somatomotor, and dorsal attention regions. Finally, edge community 5 is distinct in that its forked triads couple hippocampus and amygdala to visual occipital and sensorimotor parietal areas, as well as the dorsal attention network. Distances between cortical endpoints are widely distributed, indicating that forked triads couple spatially proximal regions within hemisphere and more distal cross hemisphere regions.

## 4 Discussion

Several studies have investigated the node-centric functional organization of cortical systems in isolation (Gordon et al., 2016; Power et al., 2011; Yeo et al., 2011) or in combination with subcortical structures such as the basal ganglia, cerebellum, and brain stem (Ji et al., 2019). The latter are especially important, since the cortex does not act in isolation, but engages in constant back-and-forth communication with subcortical regions such as the striatum, thalamus, hippocampus, and amygdala. For example, thalamic nuclei serve as information relays for sensorimotor and other information to the cortex, while also involved in cognitive function (Halassa & Kastner, 2017; Hwang, Bertolero, Liu, & Esposito, 2017; Mitchell, 2015; Sherman, 2007). Additionally, the striatum plays a role in various functions, via parallel integration of information from distributed areas of cortex (Bartoň et al., 2020; Di Martino et al., 2008; Reig & Silberberg, 2014). Finally, the amygdala is important for attribution of emotional valence (Ball et al., 2009; Jin, Zelano, Gottfried, & Mohanty, 2015) and is coupled with the hippocampus among other regions to facilitate memory encoding (Phelps, 2004; Richardson, Strange, & Dolan, 2004; A. P. R. Smith, Stephan, Rugg, & Dolan, 2006). In our recent work we have shown that edge community structure shares similarities with canonical RSNs, such that edge communities of nodes within RSNs were more similar compared to nodes between RSNs. Additionally, these edge communities coupled cortical resting state networks (RSNs) to one another, whereby multiple edge communities were identified within each canonical RSN (Faskowitz et al., 2020). Furthermore, when examining the diversity of edge community structure, heteromodal association systems (limbic, control, and default mode) showed greater diversity of edge communities compared to primary systems (visual, somatomotor, and attention) (Jo et al., 2020). Here, our aim was to understand the role of key subcortical structures (striatum, thalamus, hippocampus, and amygdala) in the edge community organization among cortical systems. Because edge communities are groups of edges with similar co-fluctuation patterns, we hypothesized that probing community organization via motif analysis can offer insight into communication patterns among node groups in a network. Therefore, we investigated the organization of edge community triads in the network that are comprised of one subcortical node and two cortical nodes, in order to identify subcortico-cortical communication patterns in the brain.

### 4.1 Functional roles of subcortical nodes defined through edge communities

Consensus clustering of data from 92 participants from the HCP100 unrelated subjects’ cohort revealed edge community structure that was related to canonical RSN organization and, within cortex, was highly similar to previously reported results (Faskowitz et al., 2020; Jo et al., 2020; Zamani Esfahlani et al., 2020) for edge communities among cortical nodes. This finding suggests that the addition of subcortical nodes to the analysis leaves cortical edge communities, which we may interpret as a proxy for intra-cortical communication, largely unchanged. The subcortex was generally partitioned into segregated communities, with its edges predominantly belonging to 3 edge communities in the 7-cluster data. The primary v. heteromodal cortical system dichotomy found in cortex-only investigation by Jo et al. (2020) and Zamani Esfahlani et al. (2020), is reinforced after the addition of subcortical nodes, where striatum, pallidum, and thalamus, and through separate communities, the hippocampus and amygdala, differentially couple to control and default mode vs. visual and somatomotor systems (Fig. 2). This is further supported by nodal entropies of cortical nodes, which were significantly different between primary and heteromodal system nodes.

There is extensive evidence for the role of core basal ganglia and thalamic regions as integrative regions that possess diverse inputs and outputs throughout the cortex (Greene et al., 2020). The edge community organization observed here, whereupon distinct communities couple the subcortex to groups of cortical systems, supports these hypotheses. These communities can be interpreted as groups of edges where the pattens of communication among connected nodes are similar to each other. With that in mind, the overlapping community structure revealed by the node-by-node matrix tells us that a node connected to some other nodes other via the same edge community has more in common with their BOLD time-courses (estimated via the co-fluctuation edge time series) compared to other nodes which connect to it by different edge communities. In that context it is plausible that these communities reveal some underlying differential coupling among nodes in a network. For instance, in our analyses the majority of subcortical nodes coupled to the cortex via 2+ edge communities, with the exception of nodes within the globus pallidus that coupled to nearly all other nodes via one community (thus resulting in entropies near zero). Additionally, while subcortical entropies tended to increase with increasing number of communities, this was not true for the globus pallidus. Assessing whether this holds true in neurological conditions, such as Parkinson’s disease, where altered globus pallidus connectivity has been reported (Miranda-Domínguez et al., 2020), may offer novel avenues for investigations into the underlying neurobiology of disease.

It is worth nothing the distinction in interpretation between edge time-series and the communities estimated from them and BOLD-dependent node-based connectivity. In the case of BOLD, regions that are said to be significantly connected will have similar nodal time-series and based on that similarity (commonly Pearson correlation) these regions are likely to end up assigned to the same community when clustering is performed. Alternatively, edge time-series capture co-fluctuation among region pairs, and clustering of edge time-series tells us which node pairs are behaving in a similar fashion (i.e., similar co-fluctuation), which can be interpreted as an indicator of similar communication patterns among brain regions. Therefore, when referring to edge communities linking the globus pallidus to other nodes in the network, grouping of its edges into a single community suggests a similar degree of co-fluctuation to other nodes, not that it is equally connected to them.

### 4.2 Distinction between primary and heteromodal systems and the role of the subcortex

Connectivity studies that utilize fMRI have tended to focus on cortical regions only for investigations of RSN structure (Power et al., 2011; Schaefer et al., 2018; Yeo et al., 2011). However, the contribution of the subcortex, cerebellum, and brainstem has gained attention in recent years (Shine et al., 2019; Tian et al., 2020), showing that subcortical network nodes also possess hub properties and contribute to rich-club organization (van den Heuvel & Sporns, 2013). This is in part due to technological, software, and methodological advancements that have allowed for better imaging and signal estimates from smaller volumes and deeper brain structures. To better understand the contributions of major subcortical structures to cortical network organization, we independently clustered edge time-series from only cortical nodes (as in Jo et al. (2020)) and from networks that included the subcortex. Our results showed a highly similar edge community structure, entropy, and similarity between the two network types. Notably, the primary and heteromodal system dichotomy reported previously (Jo et al., 2020) is still evident when the subcortex is accounted for, with subcortico-cortical interaction edges also showing this split, by coupling to the two system types via different edge communities and showing higher edge community profile similarity to primary over heteromodal systems.

These results are intuitively expected as the brain functions as a whole, regardless of whether we obtain observations from a portion of cortex, full cortex, or cortex and subcortex. That is, cortical regions receive ‘information’ *via* subcortical projections to cortex, irrespective of whether or not we actually analyze those subcortical nodes in our networks or not. While technological limitations may prevent accurate fMRI measurements of small nuclei located in the subcortex, we can continue to improve our understanding of the role of subcortical regions that we can measure. An important consideration is the relative scale at which the cortex and subcortex are measured. Here, we used a 32 node parcellation of the subcortex (Tian scale II) and a range of cortical scales from 100 to 400 nodes (Schaefer et al., 2018), so the relative ratio of cortical to subcortical nodes did not exceed ~30%. These outcomes could vary if the number of subcortical nodes added to the network was closer to equal or exceeded the number of cortical nodes. An additional consideration is the spatial resolution of human fMRI data. Subcortical regions are typically comprised of several small nuclei, which cannot be measured separately with human fMRI due to spatial resolution constraints. Insight into cortico-subcortical interaction of subcortical functional nuclei can be obtained from primate and rodent imaging data, which have shown to possess RSN structure (Belcher et al., 2013; Hori et al., 2020). Connectivity of subcortical regions is conserved, to a degree, among human, primate, and rodent species, offering an avenue for applications of edge-centric methodology in evolutionary neuroscience. Future investigations will need to be cognizant of these considerations, when assessing a more ‘complete brain network’ that could, in addition to subcortical nodes, include cerebellum and brain stem as well.

Given that the cortical edge community organization is not significantly altered when the subcortex is included, what can subcortico-cortical coupling tell us about organization of cortical brain regions? To answer this question within the framework of edge community structure, we computed entropies and edge community profile similarities for cortical and subcortical nodes, only from the edges that connect between them. Across all RSNs and subcortical regions, entropies were lower when computed from the interactions, compared to those from the full networks. The patterns of lower values for heteromodal vs. primary systems and for striatal/thalamic vs. hippocampus and amygdala were also still evident. Additionally, edge community profile similarities were nearly identical for subcortical and for cortical regions. This tells us that from the perspective of the cortex, the edge community coupling of cortical nodes within system type is similar to their coupling to the subcortex. Alternatively, from a subcortex-centric perspective two possible explanations exist: 1) subcortical nodes couple to an existing cortical organization that arises from connectivity among cortical regions, or 2) the subcortex, though its connectivity to the cortex, shapes cortical organization to some degree. Existing literature has shown that in the thalamus, different nuclei differentially couple to primary vs. heteromodal systems, with some degree of overlap to serve as an integrative hub between the two system types (Hwang et al., 2017). Clinically, disruptions in connectivity to one system type or the other, may underlie some neurological conditions, such as reported hyperconnectivity between striatum and primary systems in autism spectrum disorder (Cerliani et al., 2015).

### 4.3 Subcortico-cortical coupling via edge community triads

Recurring patterns of connectivity within networks referred to as motifs, have been employed to study organizational properties of structural and functional networks (Battiston et al., 2017; Olaf Sporns & Kötter, 2004), with an emphasis on small motifs, such as 3-node triads. While it is straightforward to quantify presence of triads in structural/sparse networks, in functional networks based on crosscorrelations, which are fully connected, topologically distinct patterns need to be derived either through thresholding or by imposing a structural backbone (Battiston et al., 2017; Morgan, Achard, Termenon, Bullmore, & Vértes, 2018). This poses a challenge as multiple thresholds must be examined to identify robust topological features. Within the edge community framework, because each edge is assigned a label, we can assess triad motif structure of a fully connected network, by examining which communities the edges of a triad belong to. This is beneficial because thresholding is no longer necessary, however, the interpretations are not the same. Assessing functional motifs on thresholded networks provides information on strength of direct connections, while edge community motif analysis examines higher order relationships (i.e., temporal co-fluctuation pattens grouped into communities). Because the purpose of this work is to probe subcortico-cortical organization, we focused on a subset of possible triads that contain one subcortical node (which we refer to as the reference node of a triad) and two cortical nodes. We found that among the four possible triad types, forked triads, where the cortical nodes connect to the subcortical node to one community while connecting to each other via a different community, were present in higher fraction than expected. In the seven-community solution, three communities had subcortical nodes with significant fraction of forked triads, grouping striatum/thalamus and hippocampus/amygdala into separate groups, with cortical triad points within either primary sensorimotor (community 1) or higher order control/default mode systems (communities 4 and 5), while attentional network connecting edge were present in all three communities that contained subcortical components.

What are the plausible interpretations for the observed triad types and the possible communication strategies underlying them? Among the four types, closed-loop and diverse triads were not significantly expressed, while L-triads were underrepresented and forked triads overrepresented. Given that we only investigated triads around subcortical nodes and only connecting to cortical nodes, this finding is consistent with our understanding of the underlying connectivity. Subcortical regions have diverse anatomical inputs and outputs throughout the cortex, which could manifest functional communication patterns of forked triads, highlighting their role in information integration and modulation of cortical activity. One must be careful not to infer directionality from these results, as such information is not available in fMRI data. It is possible that the forked triads are manifesting as a result of anatomical feedback loops between cortex and subcortex (Haber, 2016), resulting in similar communication patters of subcortico-cortical edges that are distinct from cortico-cortical communication (forked triad), and supporting synchronous zero-lag communication through mutually coupled subcortico-cortical node pairs (Gollo, Mirasso, Sporns, & Breakspear, 2014). The overrepresentation of forked motifs in nodes of both the striatum and thalamus is likely related, due to polysynaptic projections from the striatum to the cortex via the thalamus and direct cortical feedback back into the striatum, however, because we examine higher-order temporal relationships of fMRI data, we cannot discern whether direct or indirect connections drive the observed results. Nonhuman imaging in primates and rodents has shown functional activation that aligns with topographically organized structural loops between the cortex and subcortex (Haber, Kim, Mailly, & Calzavara, 2006; Han et al., 2021). Therefore, future investigations of edge community triad organization in such datasets are necessary to assess whether there is a relationship between functional network-derived edge community triads and structural subcortico-cortical loops.

Here we examined the interaction between nodes within the subcortex (striatum, thalamus, hippocampus, amygdala) and cortical RSNs as defined by Yeo et al. (2011). Choi, Yeo, and Buckner (2012) extended the cortical RSN parcellation into the subcortex by examining fMRI data and assigning subcortical regions to cortical RSNs. They showed that the striatum is connected to multiple functional systems with striatal zones connecting to district cortical RSNs and we acknowledge that there is likely integration of multiple systems in single striatal zones that was not captured with their ‘winner-take-all’ strategy. Our approach highlights that at the macros scale of anatomical regions, single subcortical nodes are coupled to multiple RSNs. That is, examining higher order relationships through clustering cofluctuation edge time series, hippocampus and amygdala showed distinct cortical coupling from the striatum and thalamus, and that all subcortical regions distinctly couple to primary vs. heteromodal systems.

### 4.4 Limitations

Limitations regarding fMRI acquisition and preprocessing must be considered when interpreting these findings. We focus on cortico-subcortical community structure and communication; however, we cannot make inferences regarding the direction of communication flow. Additionally, the edge time-series framework captures higher order relationships which may complicate the interpretability of the presented findings. Furthermore, an ongoing challenge for edge-centric analyses is to establish which higher order features might be uniquely resolvable at this scale of analysis, versus which features are accessible using static FC alone (Novelli & Razi, 2021). Such a challenge necessitates a further explorations of communication pattern dynamics, and how the topology of the structural network supports the unfolding of such patterns (Pope, Fukushima, Betzel, & Sporns, 2021). There are well understood limitations in network neuroscience regarding parcellation selection and selection/implementation of community structure algorithms. To address those were employed a multiscale cortical parcellation and performed several clustering iterations of the data with consensus clustering to ensure robustness of the present findings. Additionally, we present our findings in two datasets, consistent of separate days of HCP acquisitions in the same participants. Finally, the analyses described here focused on interactions of basal ganglia and related subcortical structures with the cortex. For a more complete brain network, future investigations can apply these methods to cortico-cerebellar and subcortico-cerebellar functional interactions. These were beyond the scope of this investigation. The cerebellum is connected to the cortex and the subcortex by multisynaptic connections via thalamus and there is evidence for thalamic integration of information coming from the striatum and cerebellum (Bostan & Strick, 2010, 2018; Hoshi, Tremblay, Féger, Carras, & Strick, 2005). To adequately assess functional roles of these connections, separate investigations, perhaps in higher spatial resolution nonhuman data, are necessary.

## 5 Conclusions

In summary, we have shown that the edge-community coupling of subcortical regions is distributed over several cortical RSN systems. As indexed by edge communities, subcortical regions differentially communicate with primary and heteromodal systems, showing greater similarity with primary systems. In a novel implementation of a triad motif analysis, prevalence of forked triads between subcortical and cortical nodes reinforces the role of the subcortex as an integrative center for information from the cortex. Future work is necessary to continue elucidating subcortical contributions in the edge time-series framework though addition of cerebellum and brainstem into functional brain networks as well as assessing whether disruptions in edge community structure offer clinically meaningful insight.

## Abbreviations

BOLD: blood oxygen level dependent;
fMRI: functional magnetic resonance imaging;
FC: functional connectivity;
rs-fMRI: resting state fMRI;
HCP: Human Connectome Project;
ETS: edge time series.

## Acknowledgement

This research was supported in-part by NSF grant no. 2023985 (R.B. and O.S.), the National Institute on Aging Training Grant on Alzheimer’s Disease and ADRD (AG071444; E.J.C), the National Science Foundation Graduate Research Fellowship under grant no. 1342962 (J.F.) and NRT grant no. 1735095 “Interdisciplinary Training in Complex Networks and Systems” (J.T., H.M., and M.P.), and the Indiana University Network Science Institute (E.J.C.). This research was supported, in part, by the Lilly Endowment, through its support for the Indiana University Pervasive Technology Institute and, in part, by the Indiana METACyt Initiative. The Indiana METACyt Initiative at Indiana University was also supported, in part, by the Lilly Endowment. Data were provided, in part, by the Human Connectome Project, WU-Minn Consortium (principal investigators: D. Van Essen and K. Ugurbil; 1U54MH091657), funded by the 16 National Institutes of Health (NIH) institutes and centers that support the NIH Blueprint for Neuroscience Research and by the McDonnell Center for Systems Neuroscience at Washington University.

## Data Availability

Imaging data come from a publicly available, human connectome project dataset, which can be accessed via https://db.humanconnectome.org/app/template/Login.vm after signing a data use agreement.

## Code Availability

Edge time-series scripts can be found at https://github.com/brain-networks/edge-centric_demo.

## CRediT Authorship Contribution Statement

**Evgeny J Chumin:** Conceptualization, Methodology, Formal analysis, Writing – original draft, Writing – review & editing. **Joshua Faskowitz:** Data Curation, Writing – original draft, Writing – review & editing. **Farnaz Zamani Esfahlani:** Writing – review & editing. **Youngheun Jo:** Writing – review & editing. **Haily Merritt:** Writing – review & editing. **Jacob Tanner:** Writing – review & editing. **Sarah Cutts:** Writing – review & editing. **Maria Pope:** Writing – review & editing. **Richard Betzel:** Conceptualization, Methodology, Software, Formal Analysis, Writing – review & editing, Funding acquisition. **Olaf Sporns:** Conceptualization, Supervision, Writing – review & editing, Funding acquisition.

---

**Supplementary Figure 1.**
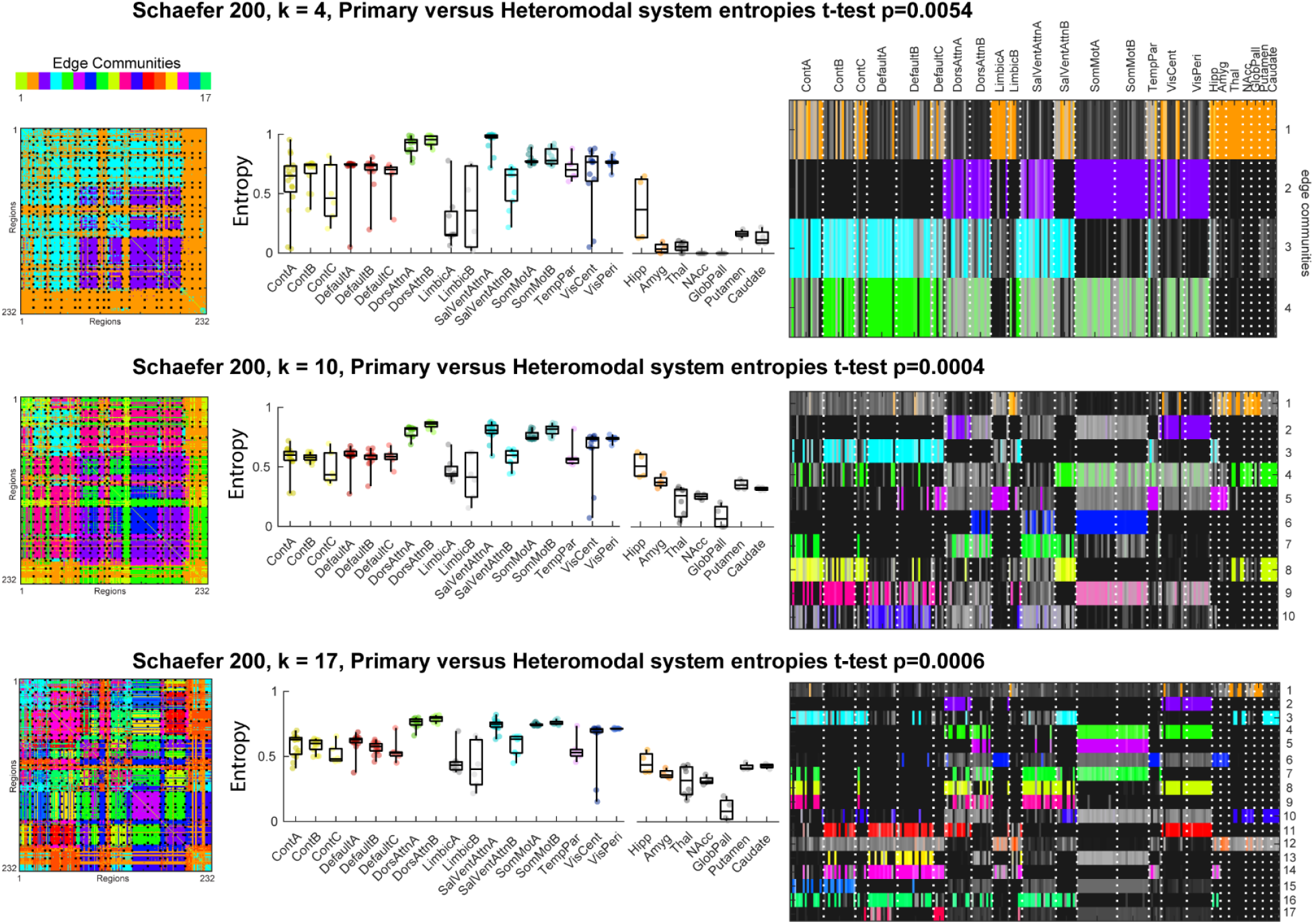
Varying number of clusters. (Left) Node-by-node matrices of edge community structure. Nodes are 200 cortical followed by 32 subcortical, ordered by 17 resting state networks (cortical) and anatomical label (subcortical). Colors indicate community labels. (Middle) Median nodal entropies of edge community structure for each of the 17 RSNs and subcortical regions, colored by 7 RSNs. (Right) Visualization of overlap in edge community structure, where color saturation corresponds to proportion of edges in an edge community (y-axis) touching a node (x-axis).

**Supplementary Figure 2.**
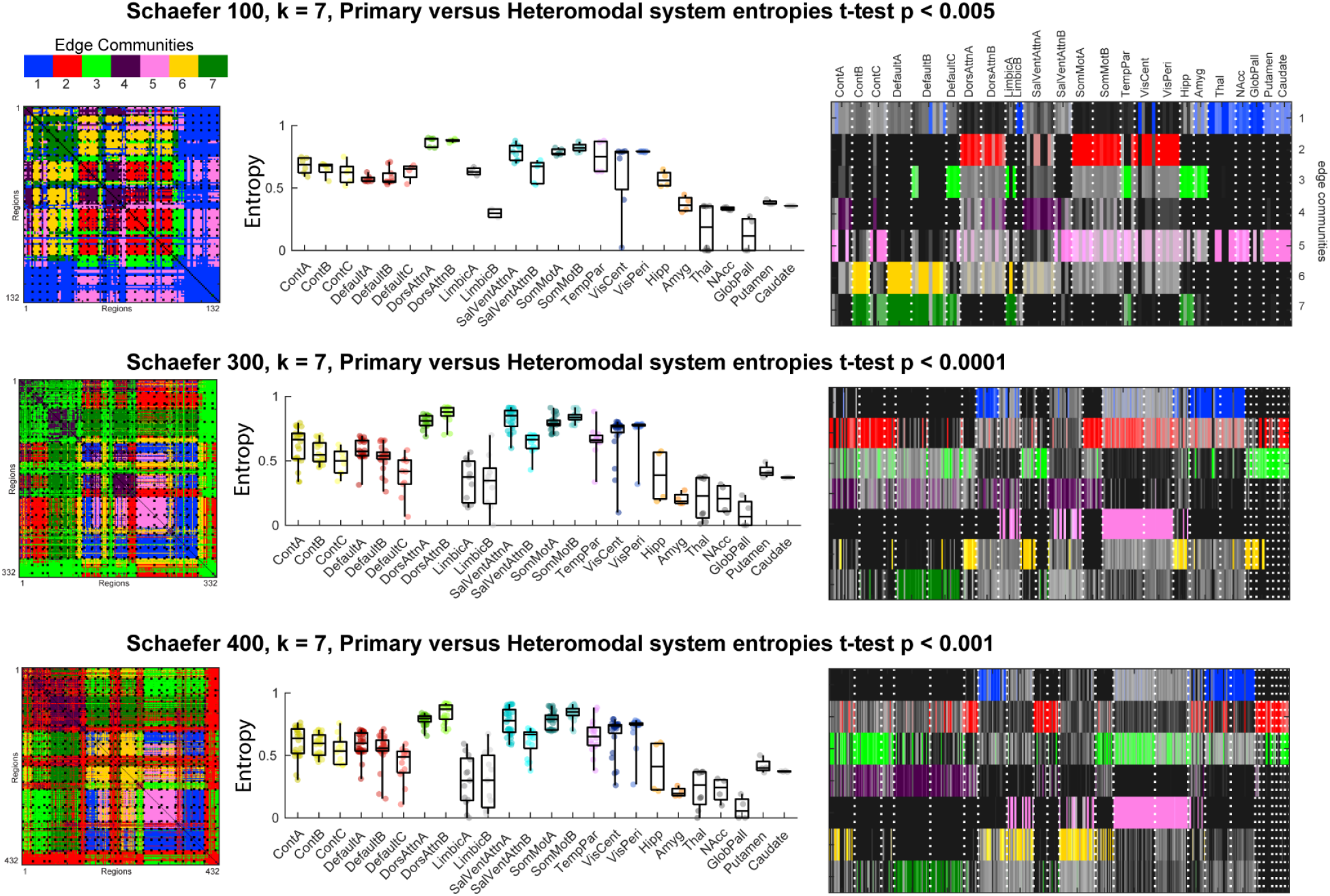
Edge community structure at various Schaefer parcellation scales with 32 subcortical nodes. (Left) Node-by-node matrices of edge community structure. Nodes are 200 cortical followed by 32 subcortical, ordered by 17 resting state networks (cortical) and anatomical label (subcortical). Colors indicate community labels. (Middle) Median nodal entropies of edge community structure for each of the 17 RSNs and subcortical regions, colored by 7 RSNs. (Right) Visualization of overlap in edge community structure, where color saturation corresponds to proportion of edges in an edge community (y-axis) touching a node (x-axis).

**Supplementary Figure 3.**
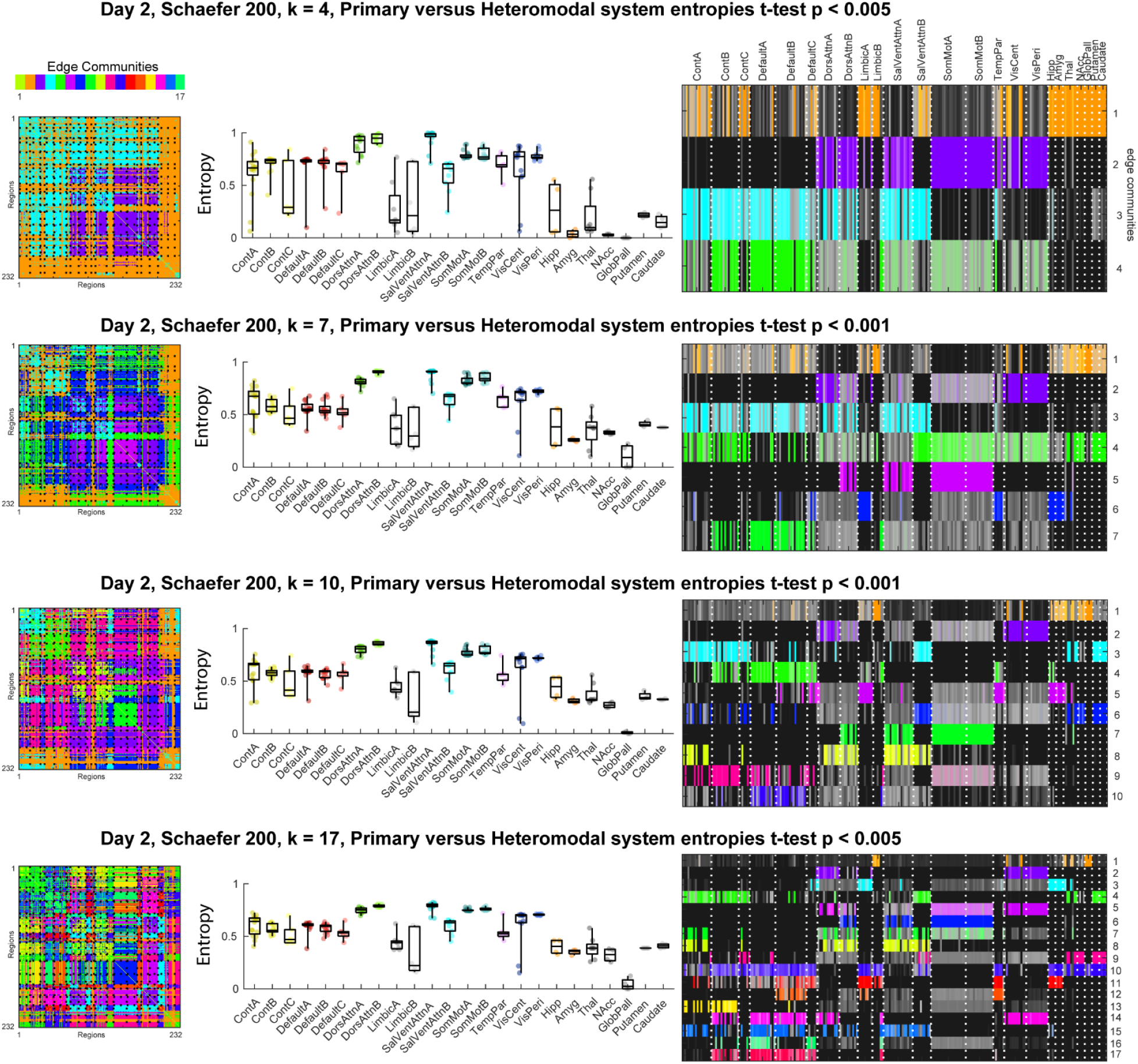
Edge community structure for Day 2 HCP data from the 92 participants in the sample at varying number of cluster/community solutions. (Left) Node-by-node matrices of edge community structure. Nodes are 200 cortical followed by 32 subcortical, ordered by 17 resting state networks (cortical) and anatomical label (subcortical). Colors indicate community labels. (Middle) Median nodal entropies of edge community structure for each of the 17 RSNs and subcortical regions, colored by 7 RSNs. (Right) Visualization of overlap in edge community structure, where color saturation corresponds to proportion of edges in an edge community (y-axis) touching a node (x-axis).

**Supplementary Figure 4.**
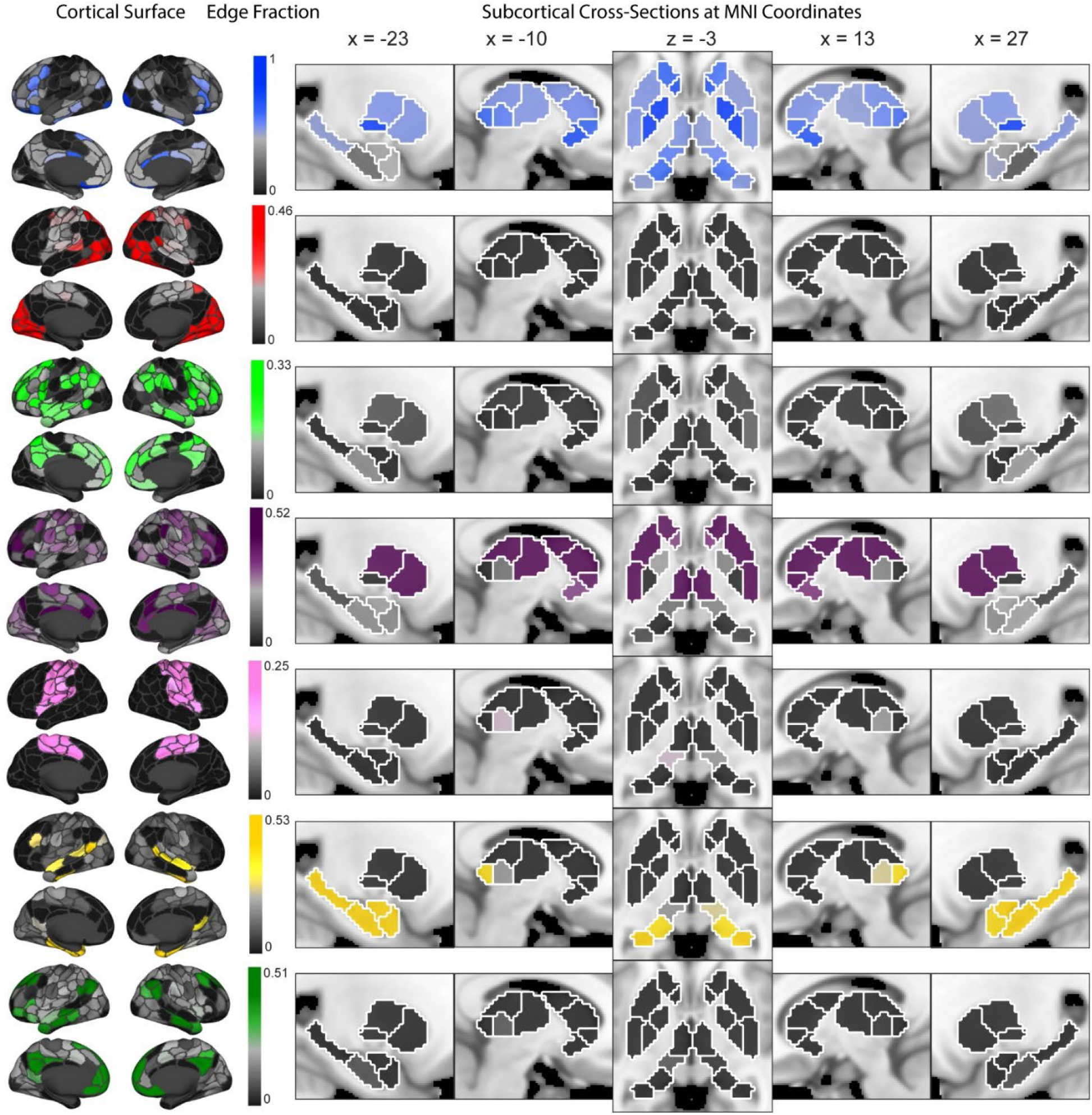
Visualization from Day2 HCP 92 participant dataset. Spatial representation of edge communities for cortical (Left; Schaefer 200 nodes) and subcortical (Right; Tian 32 nodes) parcellations. Color saturation corresponds to proportion of total edges in an edge community that connect to a particular node. Cortical nodes are visualized on a fs_LR_32k surface, while subcortical nodes are sagittal slices and an axial slice in Montreal Neurological Institute (MNI) standard space at the indicated coordinates. Anatomical underlay is the MNI152_1mm brain template. Slices left of the axial slice are in the left hemisphere while slices to the right are in the right hemisphere.

**Supplementary Figure 5.**
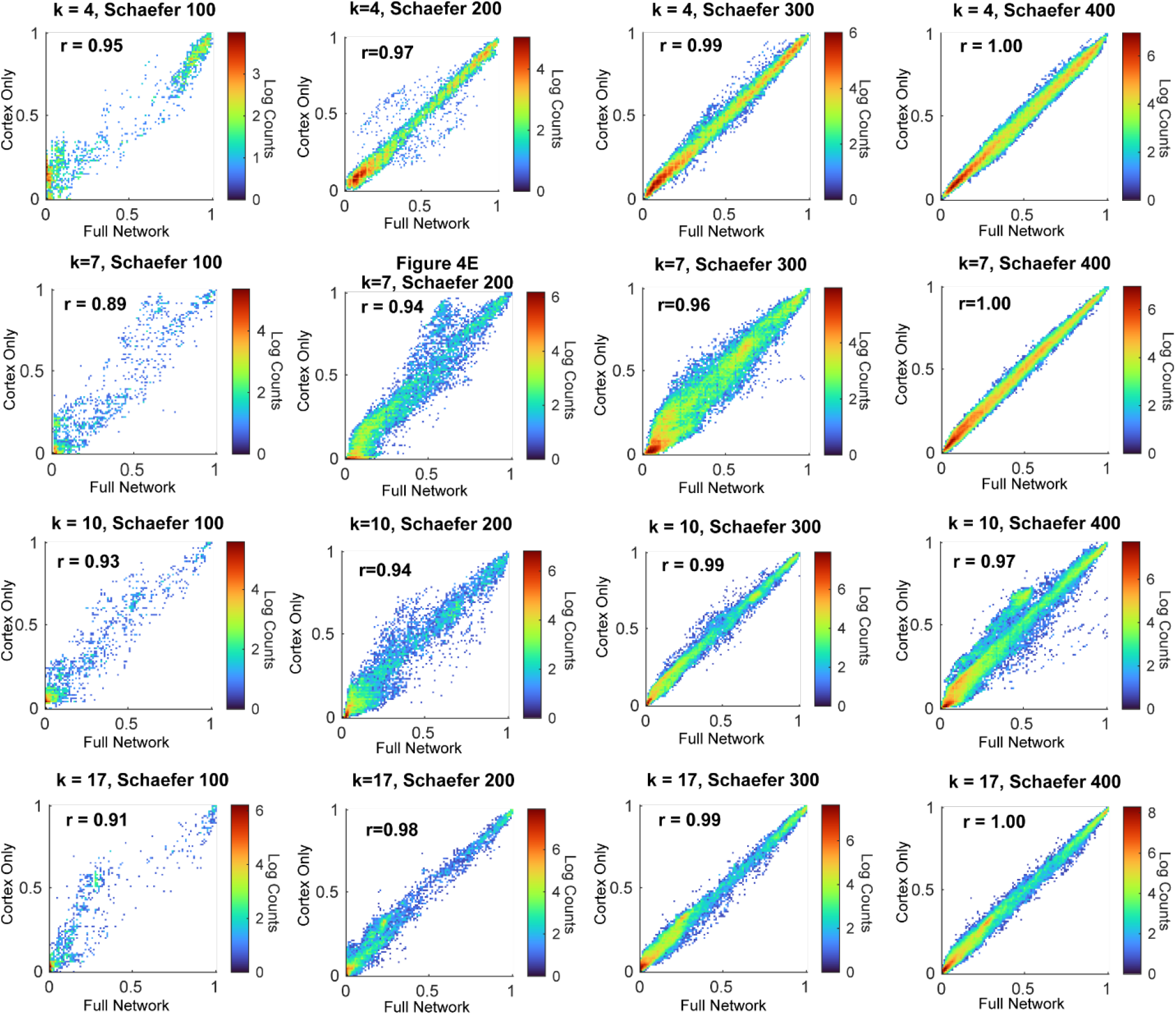
Relationships of edge community profile similarity for cortical nodes from cortex only clustered matrices vs. cortex and subcortex (full network) clustered matrices at various number of edge communities (k = 4, 7, 10, 17) and with a range of cortical parcellation scales (Schaefer 100, 200, 300, and 400 cortical nodes). For the full networks 32 subcortical (Tian scale II) nodes are present in the networks. Data are shown as 2D heatmaps on logarithmic scale. r – Pearson correlation coefficient.

**Supplementary Figure 6.**
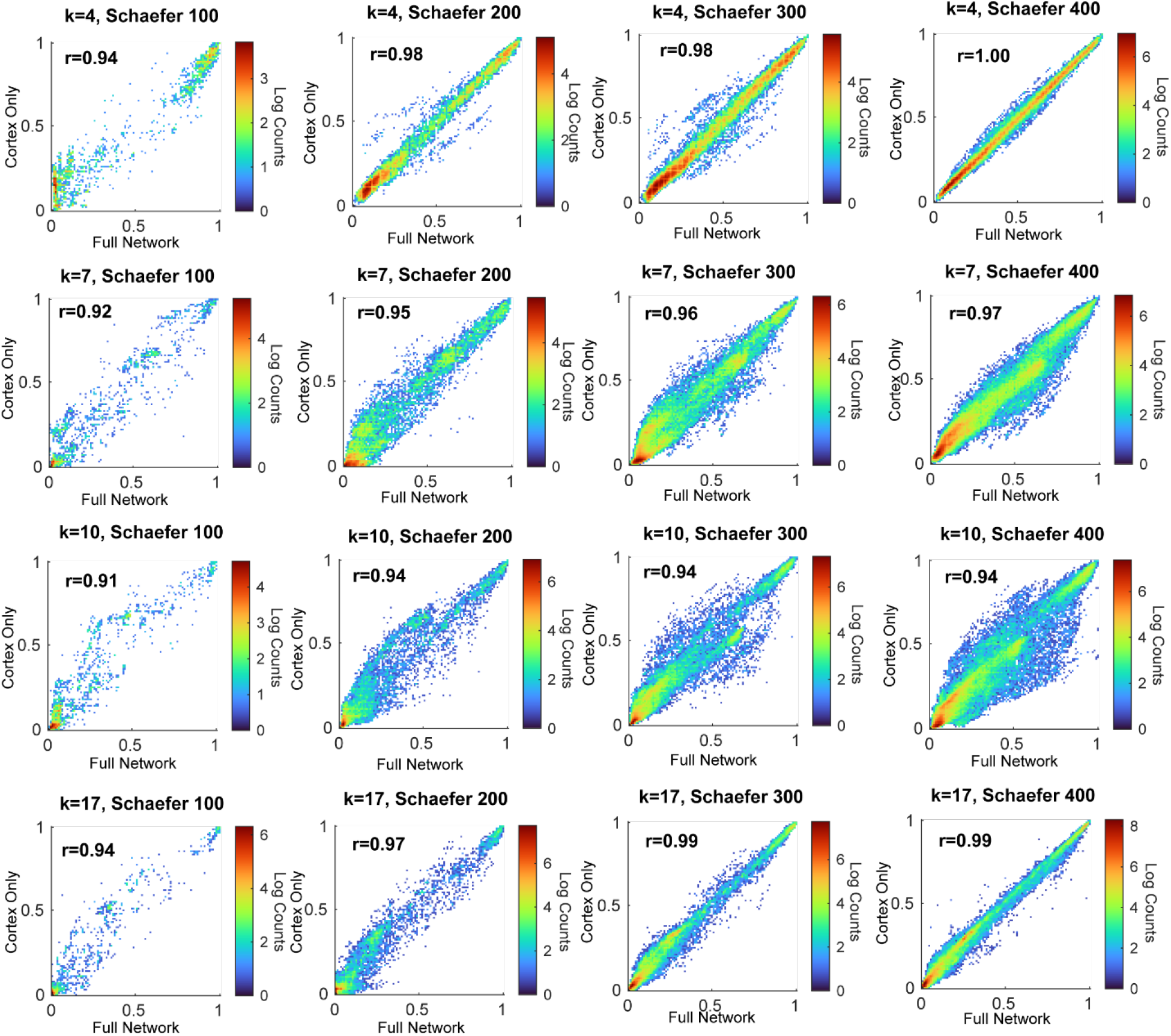
Day 2 HCP 92 subject dataset relationships of edge community profile similarity for cortical nodes from cortex only clustered matrices vs. cortex and subcortex (full network) clustered matrices at various number of edge communities (k = 4, 7, 10, 17) and with a range of cortical parcellation scales (Schaefer 100, 200, 300, and 400 cortical nodes). For the full networks 32 subcortical (Tian scale II) nodes are present in the networks. Data are shown as 2D heatmaps on logarithmic scale. r – Pearson correlation coefficient.

**Supplementary Figure 7.**
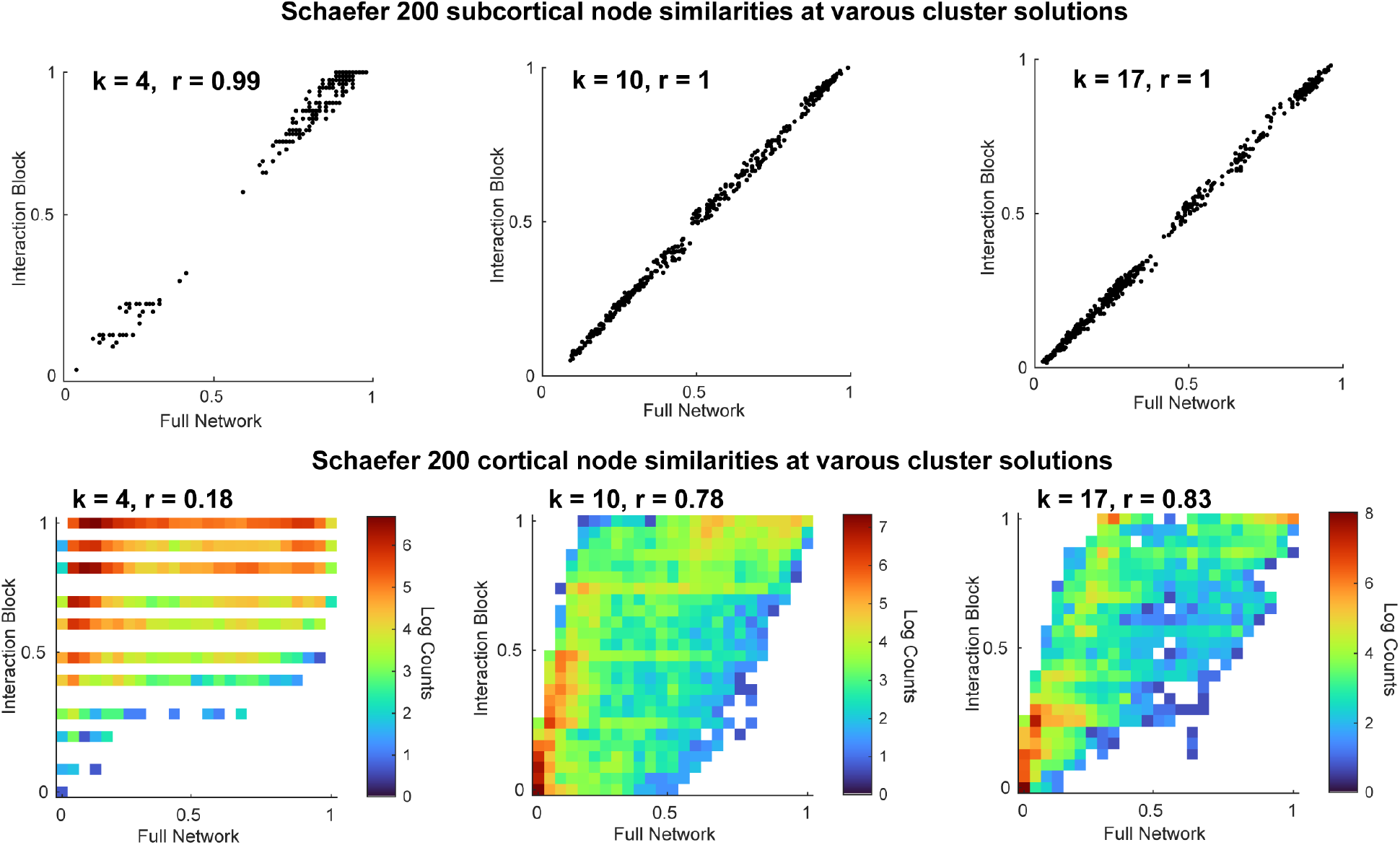
Comparisons of edge community profile similarities computed for subcortical (Top) and cortical (Bottom) nodes from the full network (x-axes) versus similarities computed only from subcortico-cortical interaction edges (y-axes) at various number of community clustering solutions. K denotes number of edge communities and r is the Pearson correlation.

**Supplementary Figure 8.**
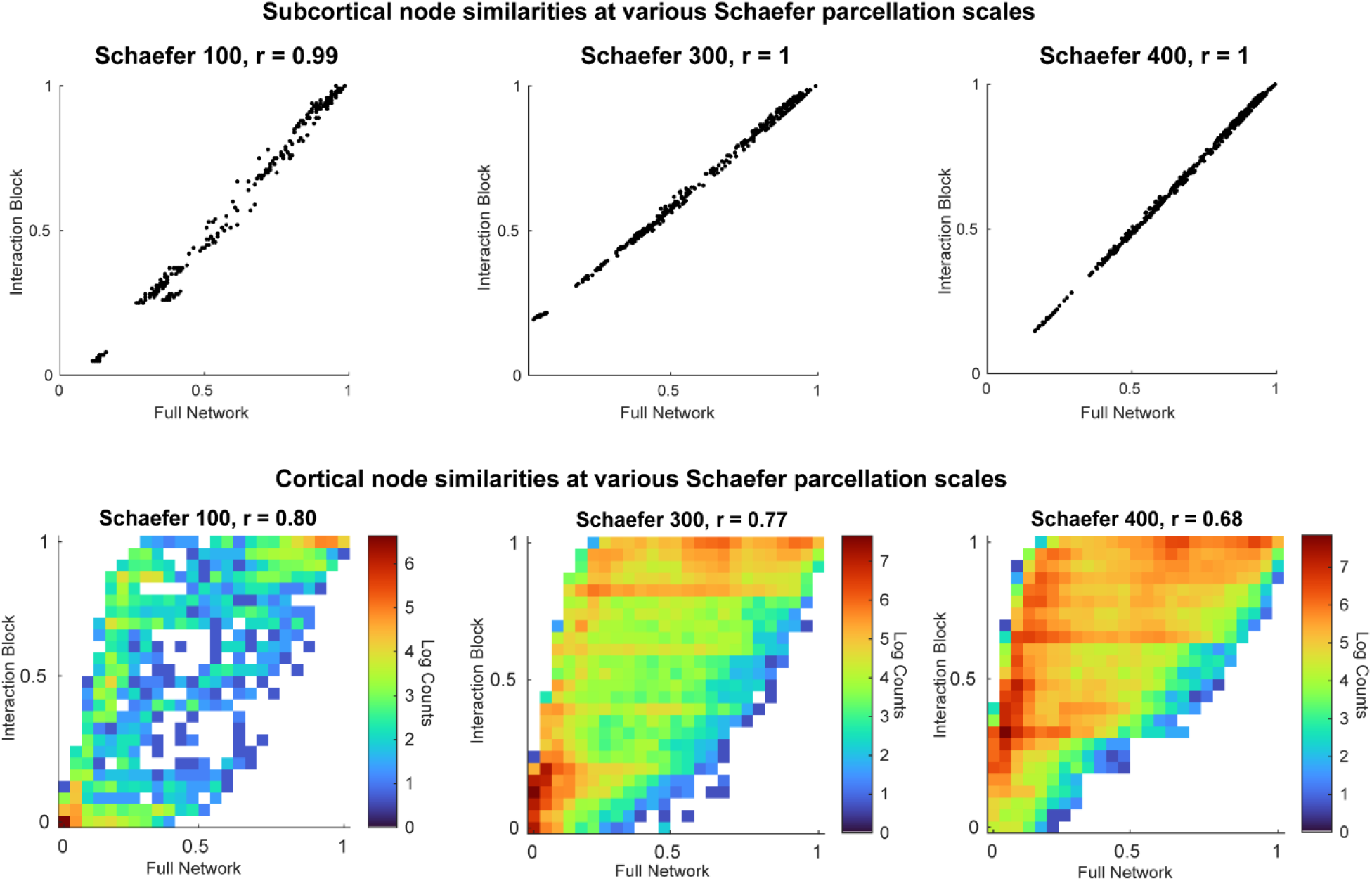
Comparisons of edge community profile similarities computed for subcortical (Top) and cortical (Bottom) nodes from the full network (x-axes) versus similarities computed only from subcortico-cortical interaction edges (y-axes) at various cortical node parcellation scales. For subcortex the Tian 32 node parcellation was used. r – Pearson correlation.

**Supplementary Figure 9.**
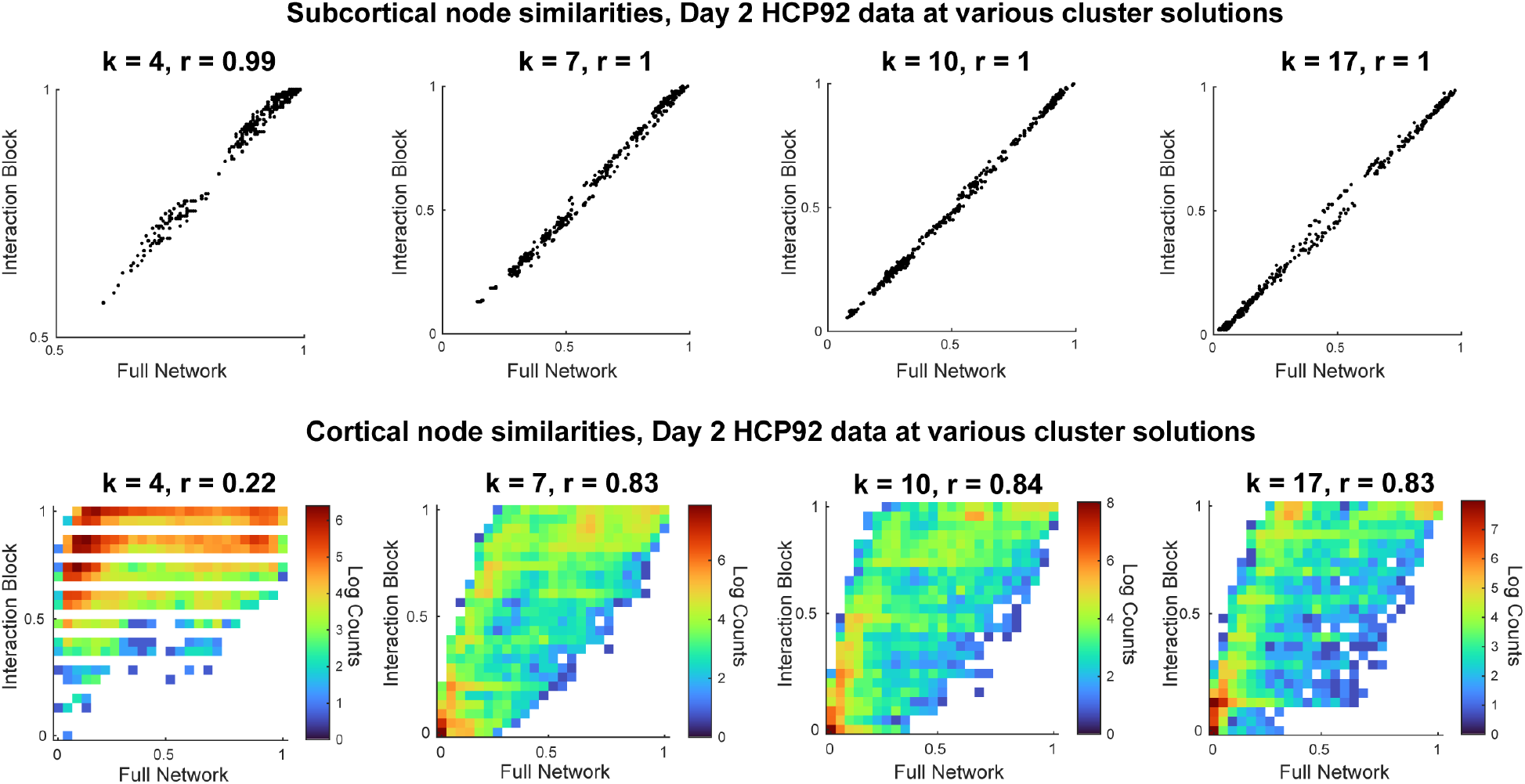
Comparisons of edge community profile similarities computed for subcortical (Top) and cortical (Bottom) nodes from the full network (x-axes) versus similarities computed only from subcortico-cortical interaction edges (y-axes) at various number of community clustering solutions in the Day 2 HCP 92 participant data. K denotes number of edge communities and r is the Pearson correlation.

**Supplementary Figure 10.**
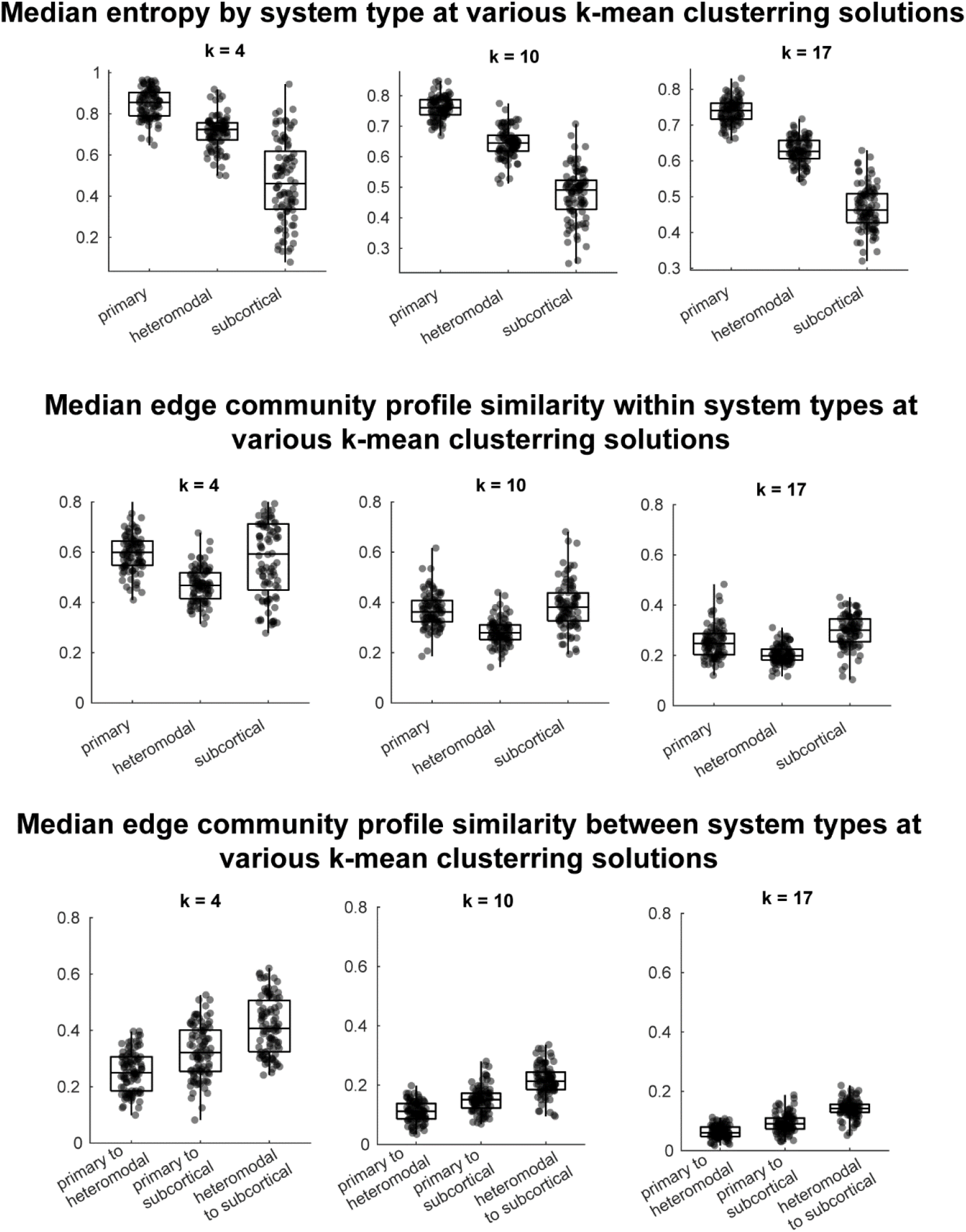
Entropy (Top) and edge community profile similarity (Middle – within, Bottom – between system types) differences between primary sensorimotor/attention and heteromodal system types, and subcortex. K denotes number of edge communities. Significant differences reported in the main manuscript (Figure 6) persisted in these data.

**Supplementary Figure 11.**
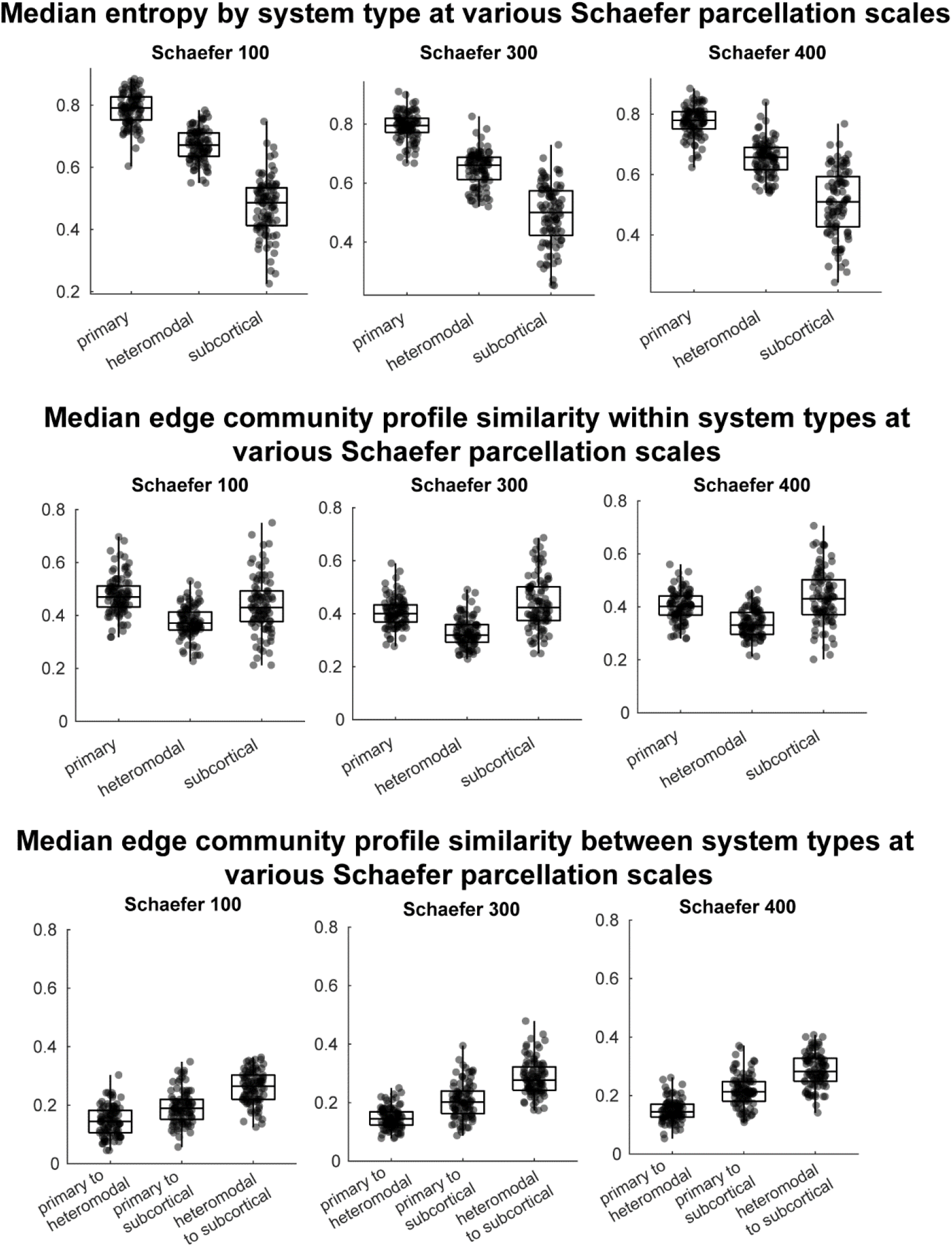
Entropy (Top) and edge community profile similarity (Middle – within, Bottom – between system types) differences between primary sensorimotor/attention and heteromodal system types, and subcortex, across a range of cortical parcellation scales. For subcortex the Tian 32 node parcellation was used. Significant differences reported in the main manuscript (Figure 6) persisted in these data.

**Supplementary Figure 12.**
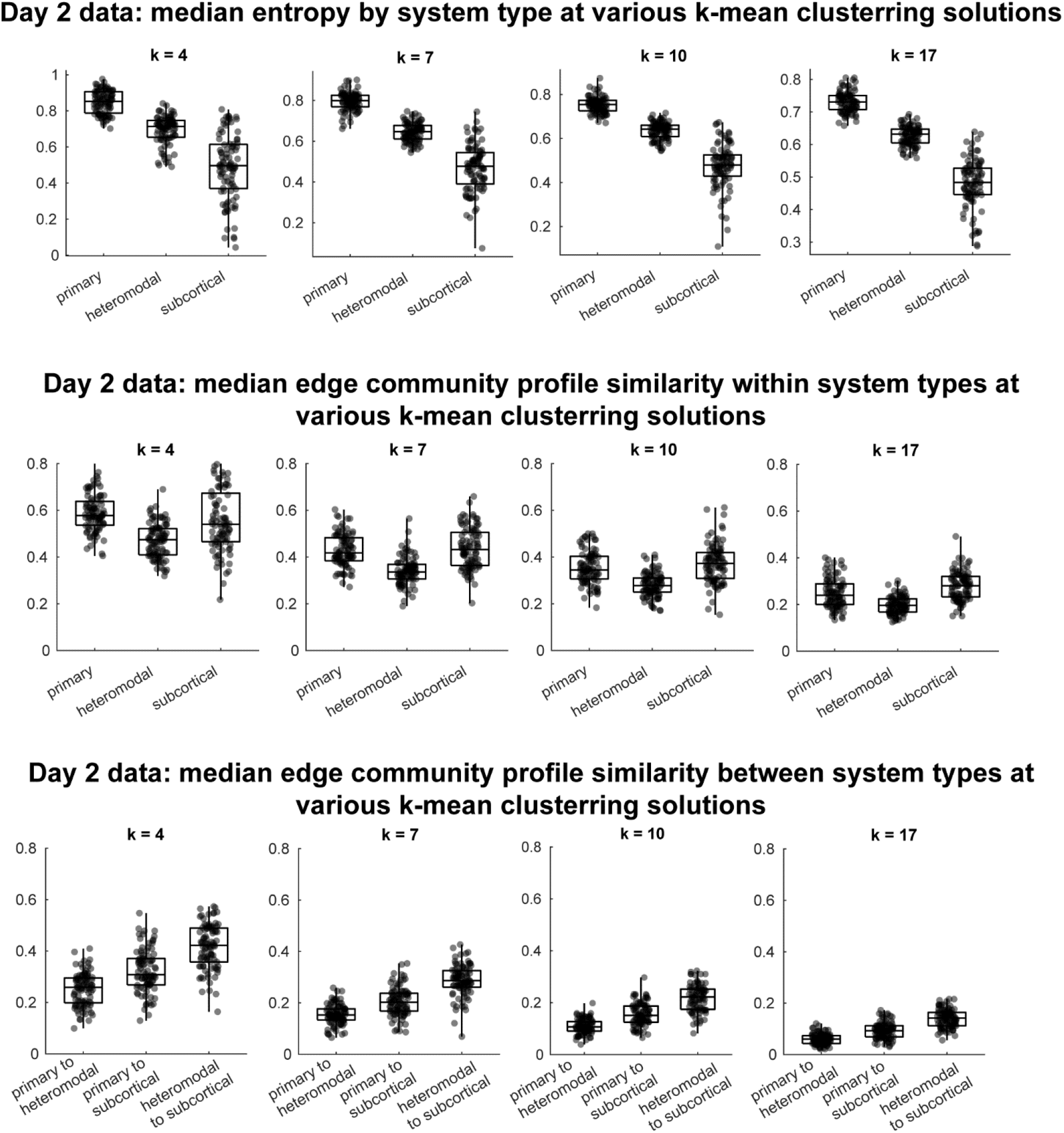
Entropy (Top) and edge community profile similarity (Middle – within, Bottom – between system types) differences between primary sensorimotor/attention and heteromodal system types, and subcortex in the Day 2 dataset from the HCP 92 unrelated participants. K denotes number of edge communities. Significant differences reported in the main manuscript for the Day 1 data (Figure 6) persisted in these data.

**Supplementary Figure 13.**
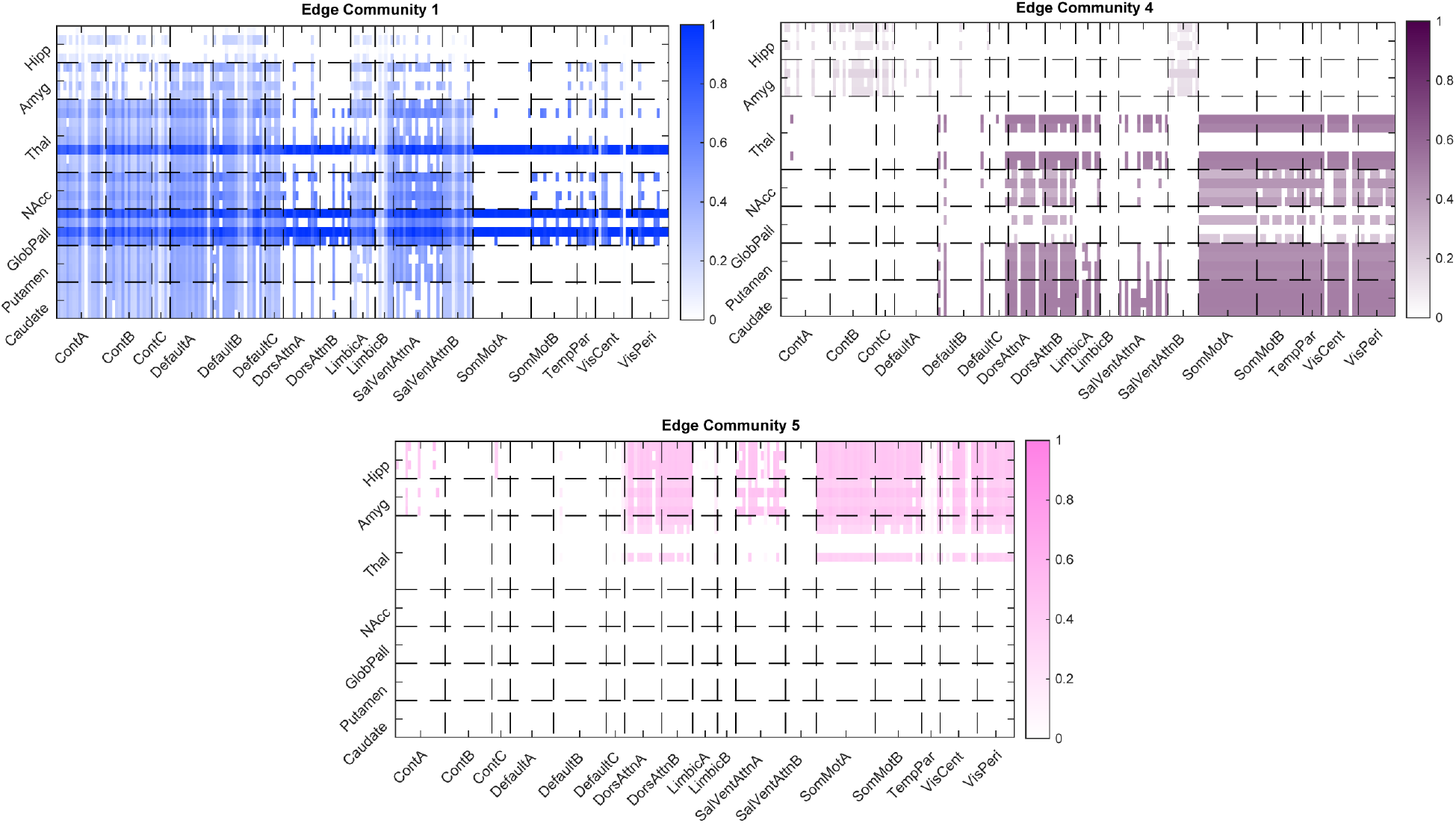
Cortical endpoints (x-axis) of triads touching one subcortical node (y-axis) for the three edge communities that showed significance in main Figure 7B. Subcortical nodes are grouped by anatomical label, while cortical nodes are grouped into the Yeo 17 RSN systems. Color saturations correspond to fraction of time a cortical node was part of a triad with a subcortical node, out of total possible triads with those two nodes. Data are from the 7-edge community matrix at Schaefer 200 node cortical parcellation scale.

**Supplementary Figure 14.**
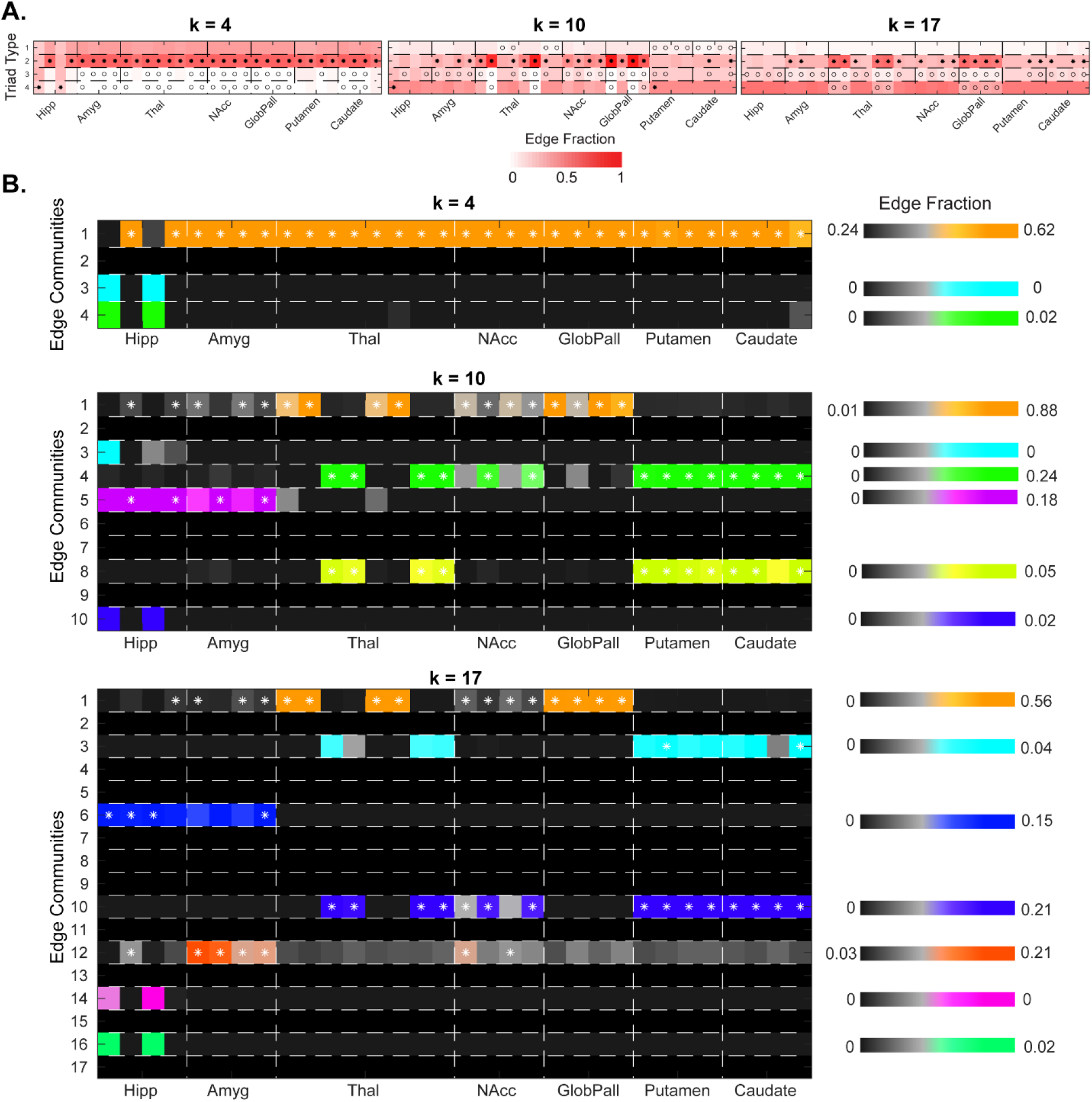
**(A)** Distributions of edge triad types at various number of edge communities. Significance was evaluated relative to 1,000 node label permuted matrices, with *p* < 0.05, two-tailed threshold, where *-denotes greater than null and o-denotes less than null. Triad types are: 1-closed loop, 2-forked, 3-L-shape, and 4-diverse (as shown in Fig. 1A). Color saturation is the fraction of total triads around that subcortical node that fall into a particular type. **(B)** A breakdown by edge community of forked-type triads, where color bars correspond to fraction of total triads around a subcortical node that are forked and fall into a particular edge community (the community that has 2 edges in the triad connecting to/from the subcortical node).

**Supplementary Figure 15.**
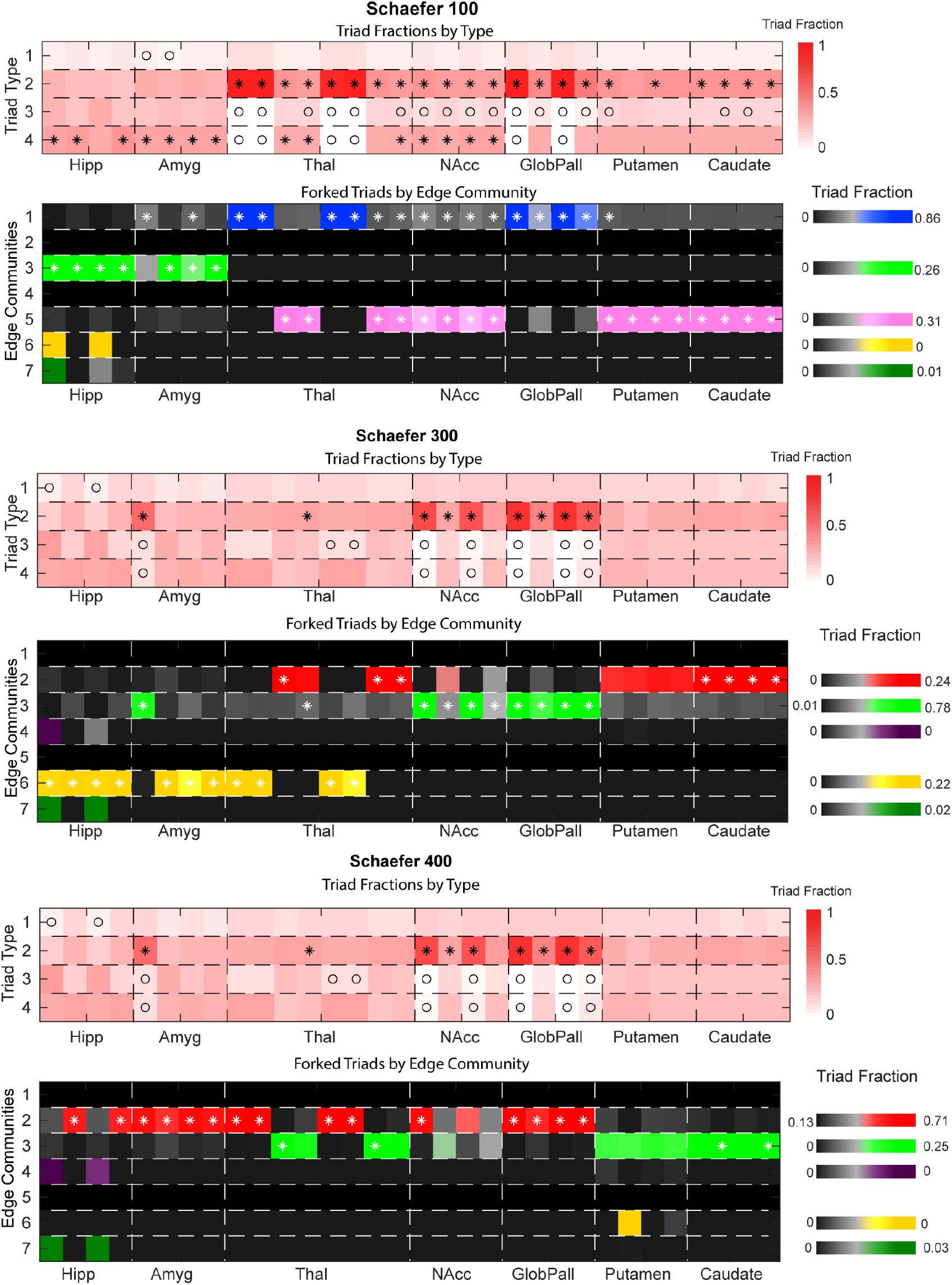
Edge triad distributions by type and for forked triads by edge community at three varying cortical parcellation scales (Schaefer 100, 300, and 400 nodes). Number of subcortical nodes was the same at all scales (Tian scale II 32 node parcellation). Color saturation

